# A computational procedure for predicting excipient effects on protein-protein affinities

**DOI:** 10.1101/2023.12.22.573113

**Authors:** Gregory L. Dignon, Ken A. Dill

## Abstract

Protein-protein interactions lie at the center of much biology and are a challenge in formulating biological drugs such as antibodies. A key to mitigating protein association is to use small molecule additives, i.e. excipients that can weaken protein-protein interactions. Here, we develop a computationally efficient model for predicting the viscosity-reducing effect of different excipient molecules by combining atomic-resolution MD simulations, binding polynomials and a thermodynamic perturbation theory. In a proof of principle, this method successfully rank orders four types of excipients known to reduce the viscosity of solutions of a particular monoclonal antibody. This approach appears useful for predicting effects of excipients on protein association and phase separation, as well as the effects of buffers on protein solutions.

**TOC Graphic:** 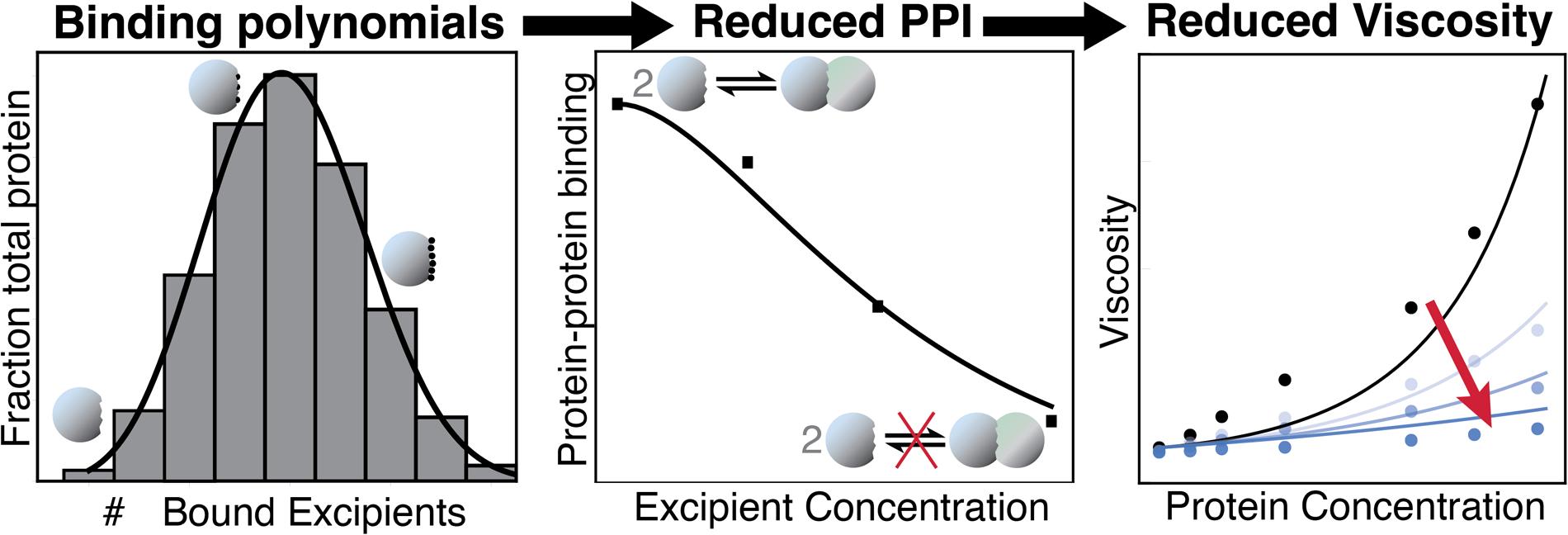

## Introduction

Proteins commonly associate into multi-protein assemblies, which can have both functional and pathological outcomes. Many multi-protein complexes such as the ribosome, and virus capsids exist in normal biology, and carry out a large number of functions. ^1,2^ More recently, much interest has been garnered by cellular membraneless organelles, which are functional assemblies of proteins and RNA molecules that act like more phase-separated polymers and less like membrane-encapsulated organelles.^3–6^ Multi-protein assemblies can also be detrimental to normal biology, such as with amyloid aggregates which are found in proteinopathies such as Alzheimer’s, Parkinson’s and more than 50 other diseases. ^7,8^ A third outcome of protein agglomeration occurs in the biologics industry, and presents a challenge in formulating biological drugs, particularly solutions of monoclonal antibodies (mAbs).^9–11^ Often, it is necessary to produce therapeutic protein drugs in concentrations that are high, but not so high as to aggregate or give high viscosity.^12,13^ Large development groups in the biotech industry seek to improve the solution properties of therapeutic protein drugs through the selection of excipients, i.e. stabilizers and salts that inhibit protein association and allow for higher concentration protein solutions to be synthesized, stored and administered. ^9,10,14–16^

In principle, molecular modeling could help with this problem by studying structure-property relationships; namely how the molecular structures of proteins and excipients lead to macroscale solution properties such as phase equilibria, viscosities, and cluster-size distributions.^17–19^ Today’s computational biology is hugely challenged to study this directly at atomic resolution, and studies that do so are limited to few systems due to the high cost and limited availability of the necessary computing resources. ^20,21^ To tackle this problem in a more high-throughput manner, it is necessary to employ a multiscale modeling approach where atomistic detail is only considered for one part of the problem. The point where atomistic detail is most necessary is in distinguishing between different excipient molecules, and their capacity to disrupt interactions between protein molecules, i.e. how does an excipient molecule disrupt 2-body protein-protein interactions? Even this is challenging at the level of atomic resolution since it is still quite computationally demanding to exhaustively sample all bound configurations of two interacting proteins, with or without excipient molecules present. Thus, it is advantageous to independently determine the bound configuration of two proteins, and then individually determine the binding of excipient molecules to the already-identified protein-binding interface on a single protein molecule,^22,23^ similar to the concept of small molecule drug design.^24^

Advances have been made on predicting excipient interactions with protein surfaces, focusing on two properties of protein-excipient binding: 1) the location of bound excipient molecules, and 2) their binding affinities.^22,23^ In this work, we hypothesize that by using binding polynomial theory, we can combine these two properties into a single parameter, thus simplifying the problem and making it possible to predict an excipient’s ability to disrupt protein-protein interactions (PPI) just from a standard MD simulation. We utilize binding polynomial theory to create what we call the *Oreo* model which gives a physical rationale for the PPI-disrupting behavior of excipient molecules. Finally, we use this information to calculate the predicted viscosity of a solution of protein and excipient based on their respective concentrations. We specifically apply this procedure to demonstrate its applicability to model data from an IgG1 mAb from MedImmune (MEDI-4212) which acheives high viscosity at moderate to high concentrations and has its viscosity lowered by various free amino acid excipients. ^25,26^

## Methods

### Modeling of protein molecules for simulations

Protein structures were retrieved from the protein databank for the Fab fragment (PDB ID: 5ANM) and the Fc fragment (PDB ID: 3AVE) of MEDI-4212. ^27^ The Fc fragment was glycosylated with a G1F glycosylation pattern using the glycoprotein builder on the glycam webserver, ^28^ at Asn-297 of each heavy chain.

### ClusPro

As a first step, we identified protein-protein binding sites by using the ClusPro webserver^29–32^ with the Balanced Energy Model. The top 10 best bound configurations were gathered and compared with experimental data to determine which regions on the Fab and Fc surfaces were most likely to participate in attractive protein-protein interactions (See Section S1; Fig. S1B,C).

### Simulation protocol

Simulations were carried out in the OpenMM 7.6 simulation engine. ^33^ Proteins were modeled using Amber ff14SB with explicit TIP3P water, ^34,35^ excipient molecules were modeled using the Generalized Amber force field (GAFF)^36^ and the glycans attached to the Fc fragment were modeled using Glycam06 force field. ^28^

The protein is initially placed in a cubic simulation box. Then a number of excipient molecules – corresponding to the desired excipient concentration – were randomly placed in the box as to not clash with protein or other excipient molecules. Finally TIP3P water was added to solvate the simulation box. These steps were carried out in Ambertools to generate the topology and input coordinate files. All systems were energy minimized, and equilibrated for 20 ns, followed by a production run of 80 ns. Simulations were conducted in the NPT ensemble using OpenMM’s Langevin Middle Integrator to keep temperature constant and propagate the system’s dynamics, and a Monte Carlo Barostat to keep pressure constant, with a 4.0 fs timestep using hydrogen mass repartitioning.^37^ Additional details for how simulations were conducted and analyzed are included in Supporting Information section S2 and Table S1.

### 7-bead antibody model and Wertheim’s thermodynamic perturbation theory

To predict the protein concentration-dependent viscosity of a protein solution, we employed the 7-bead antibody model and Wertheim’s thermodynamic perturbation theory as used previously. Briefly, antibodies are modeled as assemblies of 7 beads, where each Fab and Fc fragment is represented by two beads, and each fragment is attached to a single central bead. The mAbs are fully flexible and participate in both self-interactions and cross interactions which are accounted for in the thermodynamic perturbation theory. We employed the head-tail model for MEDI-4212 as it has been shown to participate in Fab-Fc interactions predominantly,^25^ which are treated using a square-well potential with a range (*ω*) of 0.18 nm (the length of a hydrogen bond) and a well depth of *ϵ*, which is dependent on how strongly the fragments interact. In this model, specific interactions occur only between a Fab and an Fc fragment, resulting in a polymeric system similar to hyperbranched polymers. ^38^ Using the weight distribution of antibody clusters or “polymers”, we then use colloidal theory to calculate the viscosity of the solution, which scales each cluster size by a power law. For a detailed description of this procedure refer to ref^17^ and Supporting information, section S3.

## Results

### The protein-excipient binding polynomial

In a protein assembly that contains multiple copies of a given type of protein, we assume that all protein-protein interfaces are roughly the same, so understanding the whole assembly rests on understanding a single protein pair. So, the basic process is *P* + *P* → *P*_2_ of two separate protein molecules coming together to form a dimer (Fig. 1A). To quantify the strength of interactions, we calculate the binding equilibrium coefficient, *K*_*p*_ = [*P*_2_]*/*[*P* ]^2^, where [*P* ] is the concentration of unbound proteins and [*P*_2_] is the concentration of dimers in solution.

**Figure 1.**
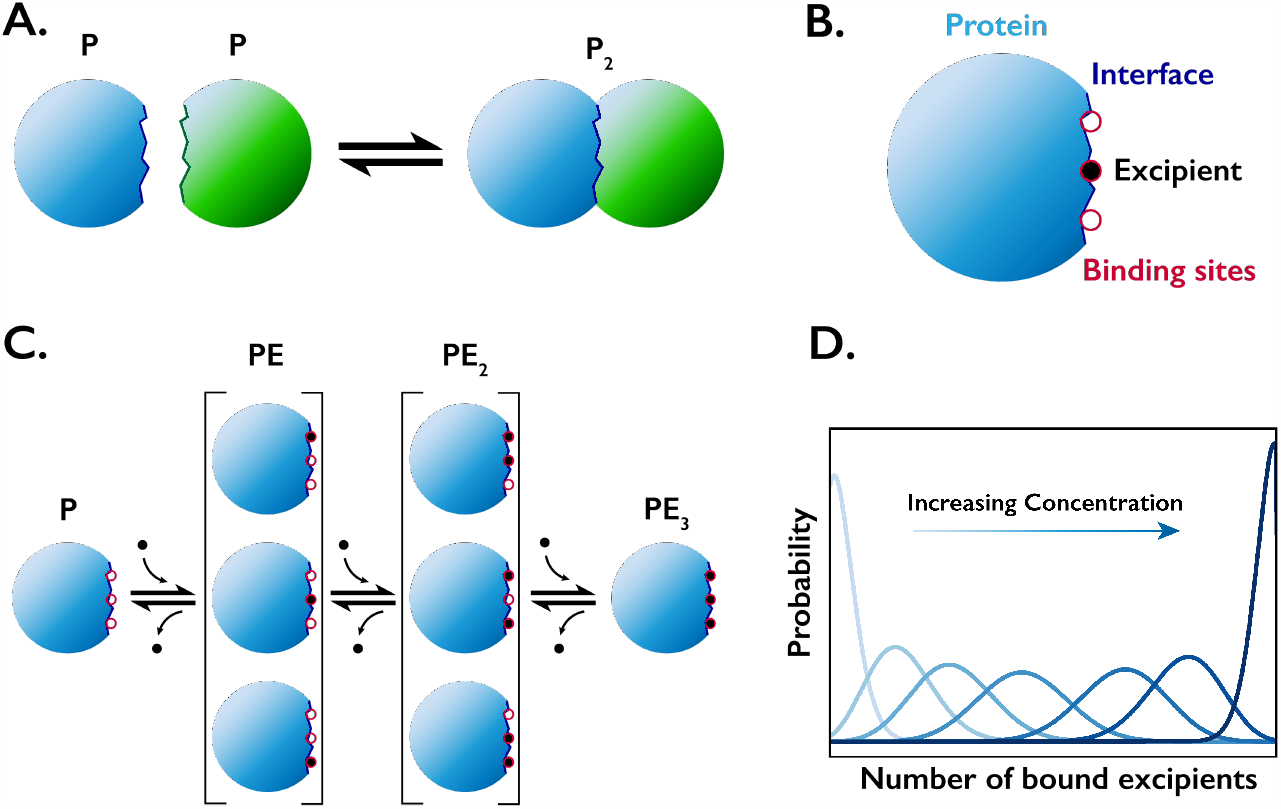
Schematic of polynomial binding model. A) Two protein molecules interact and form a dimer by fitting together at a defined interface. B) A single protein molecule has a defined interface, and several sites at which an excipient molecule may bind to. C) A single protein may bind any number, and any arrangement of excipient molecules up to the fully saturated number of binding sites, *N*. This example shows a protein-excipient pair where *N* = 3. D) Within the model, the probability of having *m* excipient molecules bound to the protein follows a binomial probability distribution. The curve shifts to the right as excipient concentration increases.

Furthermore, in order to understand how the whole protein assembly is weakened or strengthened by some added excipient in the solution (salts, small-molecule excipients or stabilizers, pH, etc.), we calculate *K*_*p*_ = *K*_*p*_(*c*), how the protein pair association depends on excipient solution concentration *c*. We do this using a binding polynomial which we derive here.^39–42^ Since excipient molecules and salts bind only weakly, we assume that all *N* excipient binding sites at the protein-protein interface are equivalent (see Fig. 1B), meaning that the average affinity of excipient for protein, *K*_*e*_, is the same for every site. Given these assumptions, the states of *m* excipient molecules binding to one of the *N* binding sites in the interface are computed from:

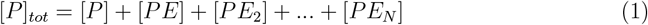

We note that this equation accounts for single protein molecules in the solution and not dimerized proteins. For each possible number of excipient molecules, *m*, up to the total number of sites, N, the concentration of the species in this series is a sum of all possible states that correspond to *m* excipient molecules. Accordingly each concentration can be expressed as a function of the number of combinations of *m* bound sites (the entropic component) and the affinity of *m* excipient molecules for a binding site (the energetic component):

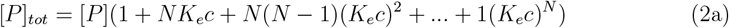

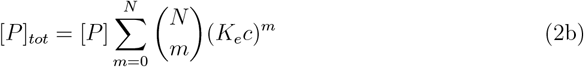

Using the binomial theorem, this can be simplified to

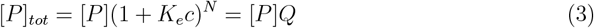

where [*P* ]_*tot*_ is the total concentration of protein in solution and *Q* = (1 + *K*_*e*_*c*)^*N*^ is the binding polynomial partition function, i.e. the sum over all possible binding states, including the unbound state. We show in Fig. 1C a cartoon schematic of a binding polynomial with *N* = 3. Within this partition function, it is notable that higher-m species acheive greater population at higher values of c (Fig. 1D). Importantly, the concentration of protein with *m* excipient molecules bound to it is given by

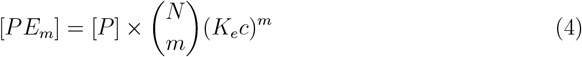

To avoid computation of the free protein concentration [P] which is non-trivial, we divide both sides by [P]_*tot*_ to, and using the equivalence from eq. 3 obtain the probability distribution function:

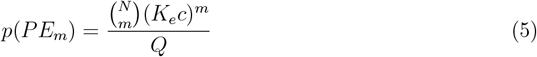

This framework may be generalized to account for excipient-binding sites that have different affinities (See SI section S4). This approach mirrors that of Aune and Tanford’s work on studying denaturation of lysozyme using guanidine hydrochloride.^39^

### The *Oreo Model* of binding interfaces

In contrast to the problem of protein denaturation where no denaturant molecules are accommodated within the folded protein structure, the protein-protein interactions we are studying may not be so strict. We must consider how many excipient molecules *m* will be allowed within the protein-protein interface. While it is difficult to tell in advance, and likely dependent on the protein interface, and excipient molecule identity, we find that three possible cases are sufficient to fit the data herein. Specifically, we develop and test three specific versions of the *Oreo Model* (Fig 2) which come from a generalized version which we will also derive here. The Oreo is a type of cookie in which two crackers (i.e., the two protein molecules) are separated by a layer of filling (i.e., the excipient molecule(s) between the two proteins). In the simplest case, the *Empty Oreo* model, the protein-protein interface contains zero excipient molecules, so the binding affinity is

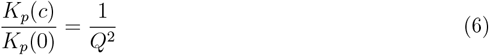

**Figure 2.**
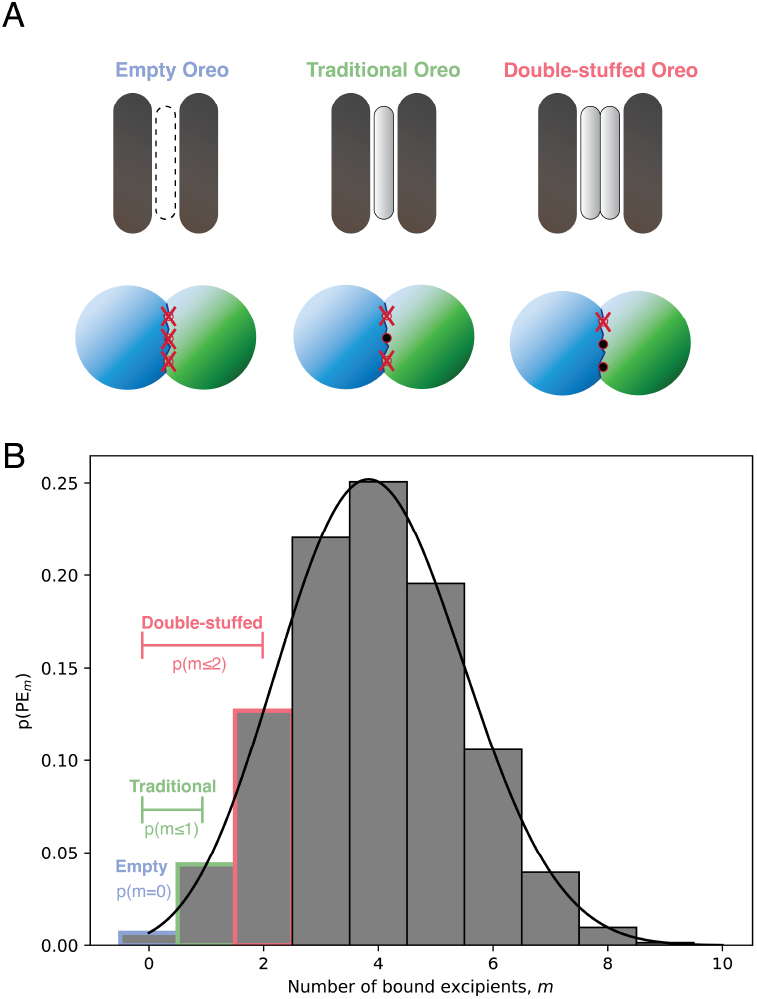
The *Oreo* model describes the number of molecules allowed within a bound configuration. A) The *Empty Oreo* model allows no excipient molecules between the two proteins, while the *Traditional Oreo* and *Double-stuffed Oreo* models allow one or two excipient molecules. B) A binomial distribution describes the population of protein molecules with *m* excipient molecules bound. The different *Oreo* models consider a different number of states as “reactive” species which may bind to other protein molecules. In this case, the *Empty Oreo* model has a probability density of about 0.01, the *Traditional Oreo* has about 0.05, and the *Double-stuffed Oreo* has about 0.18.

The denominator in Eq 6, *Q*^2^, represents the 2 isolated proteins before the protein-protein binding event. The numerator, 1, represents the singular state in which no excipient is bound to either protein interface, which occurs when bound proteins squeeze out all possible excipient molecules. Thus, we are effectively modeling competitive binding between an ensemble of possible combinations of excipient molecules, and a single protein binding partner on the PPI interface.

It is not unreasonable, however, to assume there may be some excipient molecule(s) accommodated within the bound interface between the protein molecules. An additional parameter, *m*_*i*_, will describe the number of excipient molecules accommodated within the binding interface, and will define which *Oreo Model* is being used. This gives rise to the *Traditional Oreo* model, and the *Double-Stuf Oreo* model (Fig. 2).

We first consider a protein-protein interface having 1 excipient bound inside (*m*_*i*_ = 1; the *Traditional Oreo* model). The binding affinity is

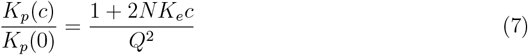

where the value in the numerator comes from the binomial distribution of how *m* excipient molecules distribute across 2*N* binding sites. Then we consider a protein-protein interface having 2 excipient molecules bound inside (*m*_*i*_ = 2; the *Double-stuffed Oreo* model). The binding affinity is

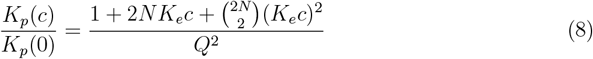

We can, of course, extend to larger values of *m*_*i*_ with a generalized *Oreo* model

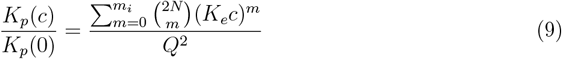

but find that results do not improve for the data studied in this work (Fig. S3). This may, however be applicable in other scenarios, perhaps with more flexibility in the protein-protein interface, or very small excipient molecules.

Thus, in order to compute *K*_*p*_(*c*) we need to know excipient affinity for the protein site, *K*_*e*_, the number of excipient binding sites on the protein interface *N*, and the number of excipient molecules sandwiched in between the bound state of the proteins, *m*_*i*_. The following section describes how we extract these quantities from atomistic computer simulations.

### Obtaining the microscopic parameters from atomistic simulations

To demonstrate the applicability of this approach, we study a single mAb, MEDI-4212 which achieves high viscosity at moderate concentrations (∼50 mg/mL), and has been found to have reduced viscosity in the presence of four free amino acid excipients (Arg, Asp, Glu and Lys).^25,26^ This mAb is known to self-associate through Fab-Fc interactions, meaning that the majority of intermolecular interactions leading to high viscosity are likely heterotypic, involving two different fragments. The binding polynomial theory and the *Oreo* models can be easily reformulated to account for heterotypic binding, which we derive in a later section.

We first identified the protein-protein interface by using the rigid protein docking web-server ClusPro^29^ to find stable bound conformations between the Fab and Fc fragments of MEDI-4212. From this, we identified two bound conformations that matched well with experimental hydrogen exchange data^25^ (Fig. S1B,C). We then defined the PPI interface explicitly as any amino acid residue that is found to be buried. The definition of the binding site is dependent on which conformation we use as well as the definition of a “buried” residue. See SI section S5 for more details (Fig. S4-6; Table S2-4). Further work on more protein-excipient systems will be beneficial in optimizing these definitions.

We also tested AlphaFold Multimer ^43,44^ as a possible alternative to finding bound configurations. Interestingly, AlphaFold Multimer did not identify bound configurations that agree with the experimental data. We suspect this may be due to the Fab-Fc interactions being relatively weak and nonspecific, which may not be captured well by algorithms that are trained on data from the protein databank, which would mostly capture strongly bound complexes.

Once the PPI interface had been identified, we used MD simulations to quantify excipient interactions with each fragment’s interaction-prone region. We simulated each fragment individually in the presence of 25, 50 or 75 excipient molecules, corresponding to concentrations (c) of 70, 140 and 210 mM with explicit solvent and ions to neutralize the system charge. If we account for the volume displaced by the large protein molecule, the effective excipient concentrations become 77, 154 and 231 mM. Each simulation was conducted for 100 ns, and the first 20 ns were discarded as equilibration time. For a full list of simulations used in this work, see Table S1.

We calculate the average number of excipient molecules bound to the protein surface at the PPI interface for both Fab and Fc, and with each excipient molecule type. We show the probability distribution of the number of excipient molecules bound to the Fc PPI interface in Fig. 3 and fit the expected binomial distributions at the three tested concentrations. Fitting these curves gives the two quantities we seek, *K*_*e*_ and *N*. We find that for most cases, there is not a unique solution to the binomial fit, but rather a range of values that fit these data well (Fig. S7). As a practical matter, we define a proxy parameter, *a* = *NK*_*e*_ which has a unique solution, and fit the binomial curve using *a* and *N* as free parameters. An advantage to using this *a* parameter is that it describes both how tightly an excipient will bind to the PPI interface, and how many excipient molecules can bind. In physical terms, the quantity *a* represents how much enhancement of concentration of excipient the whole protein (or surface region) provides relative to an equivalent amount of bulk volume, and statistically, it is related to the maximum value of the probability distribution.

**Figure 3.**
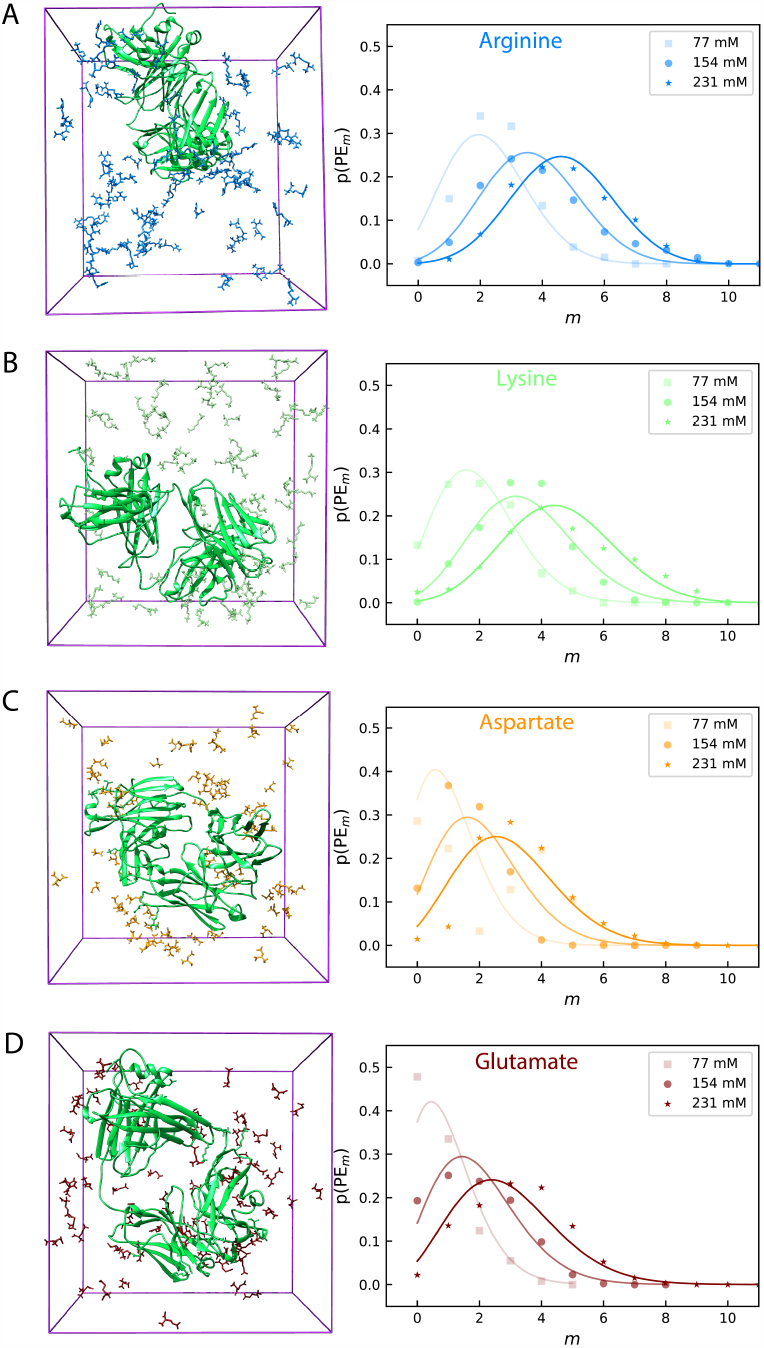
Simulation snapshots and binding polynomials to the PPI interface of the Fab fragment. Simulation snapshots show protein in green and excipients A) arginine, B) lysine, C) aspartate, and D) glutamate. Symbols in plots show the histogram of the number of excipient molecules bound to the PPI interface identified by ClusPro model 3. A binomial distribution was fit to histograms from three different simulations at different excipient concentrations. Fits are shown as solid lines, and the fitted parameters (*N*_*B*_ and *k*_*E*_) are shown at the top of each figure.

The values of the parameters *N, K*_*e*_, and *a* that we find from our MD procedure above are given in Table 1, and the values of *a* for different excipients shown in Fig 4. Of the four excipients we tested here, the results show that arginine is predicted to be the strongest disrupter of these protein dimers and glutamate is the weakest, which appears to be the case from the experimental studies too and will be explored in more detail in the following section.

**Table 1:** Optimized parameters for binomial fits to Simulation data. Bounds were set for the *N* between 10 and 100 since the optimal solution is not very sensitive to the value of *N* (See SI section S5).

|  | Fab |  |  | Fc |  |  |
| --- | --- | --- | --- | --- | --- | --- |
| | $N$ | $K_e$ | $a$ | $N$ | $K_e$ | $a$ |
| Arg. | 10.0 | 3.707 | 37.075 | 10.0 | 1.500 | 15.002 |
| Lys. | 13.9 | 2.102 | 29.266 | 10.0 | 1.408 | 14.083 |
| Asp. | 24.6 | 0.581 | 14.264 | 10.0 | 0.740 | 7.396 |
| Glu. | 100.0 | 0.127 | 12.703 | 10.0 | 1.129 | 11.291 |

**Figure 4.**
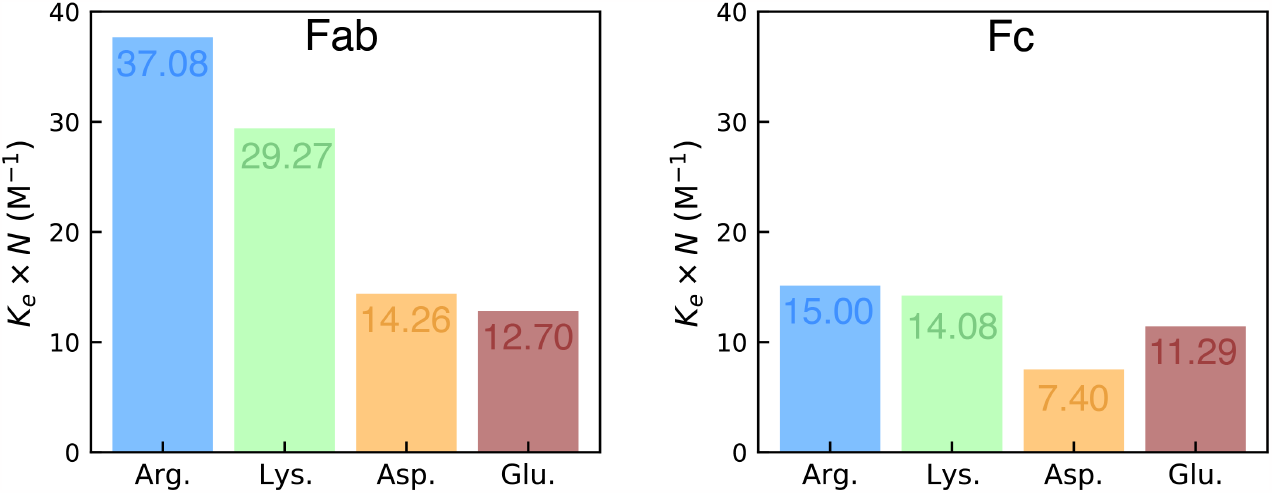
Calculated excipient quality from simulations, *a* = *K*_*e*_ × *N*, for each excipient molecule tested. These parameters were fit using the PPI interface defined by a 4°*A*^2^ cutoff for ClusPro model 3.

### From the microscopics to macroscopic predictions

The simulations above provide us two parameters that we need for computing the protein dimerization equilibrium coefficient *K*_*p*_(*c*) as a function of the excipient concentration, from a general-use atomistic model. We now seek two additional insights: (1) Is this dimerization affinity consistent with macroscopic properties of whole aggregates, such as the solution viscosity, liquid-liquid phase equilibria, and cluster size distributions? And (2) Which *Oreo* model is most appropriate to use for this protein-protein interface?

To address the first of these questions, we employed Wertheim’s thermodynamic perturbation theory as in previous work^17^ to calculate viscosity curves from site-site interaction strength of antibody molecules. We employed the head-tail model for MEDI-4212 as it has been shown to participate in Fab-Fc interactions predominantly.^25^ Thus, specific interactions occur only between a Fab and an Fc fragment, resulting in a polymeric system similar to hyperbranched polymers. ^38^

Since it isn’t completely straightforward to determine which Oreo model will be most appropriate, we first need an estimate of how strong the protein interactions must be, and how they are affected by excipients. Therefore, we employ the Wertheim theory in reverse, using the experimental viscosity curves as a starting point, and calculating the binding strength (*ϵ*) parameters from the head-tail model. We find that the Wertheim model fits the viscosity data from the experimental papers well (Fig. 5A, S8). Using the Wertheim model, we can also convert this value to an association constant, *K*_*p*_ by comparing the number distribution of monomers and dimers at very low concentration.

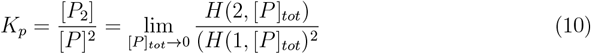

where H is the number distribution of protein molecules in the solution. The parameters *ϵ* and *K*_*p*_ follow the thermodynamic relationship

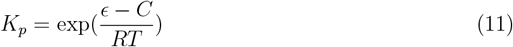

where C is equal to 18.2 kJ/mol and is related to the loss of entropy upon dimerization. The results for *ϵ* and *RT* ln(*K*_*p*_) for each of the experimental viscosity curves are included in Fig. 5B, and the linear fit between the two is shown in Fig. S9.

**Figure 5.**
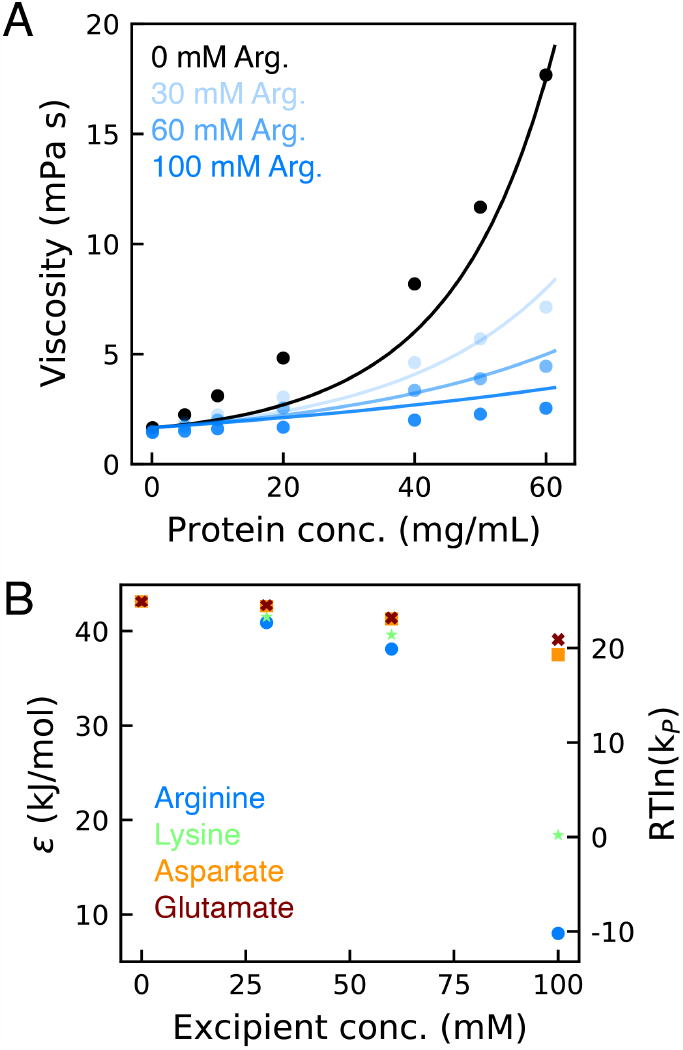
A) Viscosity curves of MEDI-4212 in solution with four tested excipients. Points show experimental data from ref. ^26^ Solid lines show fit using Wertheim theory. B) Values of binding energy and association constant as a function of excipient identity and concentration. Values were extracted from fitting the Wertheim theory to experimental viscosity curves.

We note that the zero-excipient case yields an *ϵ* value of 43.2 kJ/mol (∼ 17.4 kT) and a *K*_*p*_ of 24,910 *M*^*−*1^, and all cases containing excipient have smaller values of *K*_*p*_. Considering there is still buffer present in all experiments, this is not a true “zero-excipient” condition as there is a small amount of histidine in the solution, about 10 mM. Since this buffer is present at a constant concentration in each of thee experiments, we can absorb it into the value of *K*_*p*_(0). To demonstrate that this is an appropriate assumption, we derive a multi-excipient version of the theory (See Supporting Info Section S6).

### Evaluating the Oreo Models

Using the information obtained from the previous sections, we may now assess how well the different *Oreo* models perform. To directly compare, we can use the values of the proxy parameter *a* calculated from simulations, and the same parameter calculated from fitting the excipient concentration-dependent association constants (*K*_*p*_(*c*)) derived from experiments in the previous section. A few important modifications need to be made to the *Oreo* models. We first must account for the heterotypic interface of MEDI-4212 which interacts through Fab-Fc interactions. We modify the generalized *Oreo* model (eq. 9) to reflect the presence of two distinct interacting surfaces, each with its own unique surface chemistry, (i.e. N, and *K*_*e*_ parameters).

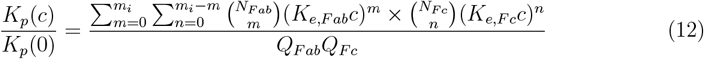

where 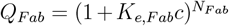and 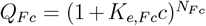. Secondly, we must substitute the proxy parameter *a* into the equation to replace either *N* or *K*_*e*_. Practically, it is easier to replace all instances of *K*_*e*_ with *a/N*. Thus the final equation we use to fit the different *Oreo* models is:

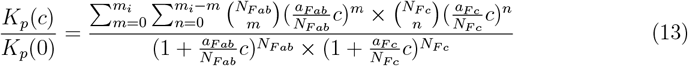

From this construction of the model, we have four free parameters that must be fit to fully describe the molecular details of the protein-protein, protein-excipient competition, *N*_*F c*_, *N*_*F ab*_, *a*_*F c*_ and *a*_*F ab*_. For convenience, we can take advantage of the definition of the *a* parameters and show that *K*_*p*_ is mostly insensitive to the values of *N*_*F c*_ and *N*_*F ab*_ (See Fig. S10A). This is because *K*_*e*_ will change to offset the effect of *N* as long as *a* is held constant. We also note that the heterotypic model can be approximated very well by the homotypic model by taking the average of the *a* and *N* values (Fig. S10B). Given these caveats, we can simplify the system down to fitting a single parameter: *a*_*avg*_ = *mean*(*a*_*F ab*_, *a*_*F c*_) which can obtained both from fitting the experimental data, and from the simulations.

In Fig. 6A-C we show the three Oreo models fit to the *K*_*p*_ valules derived from the experimental data. While all three models seem to fit the data reasonably well, the values of the fit parameter *a*_*avg*_ are quite different between the three models. Indeed, the Traditional Oreo model has the best quantitative agreement with the *a*_*avg*_ parameters obtained from simulations.

**Figure 6.**
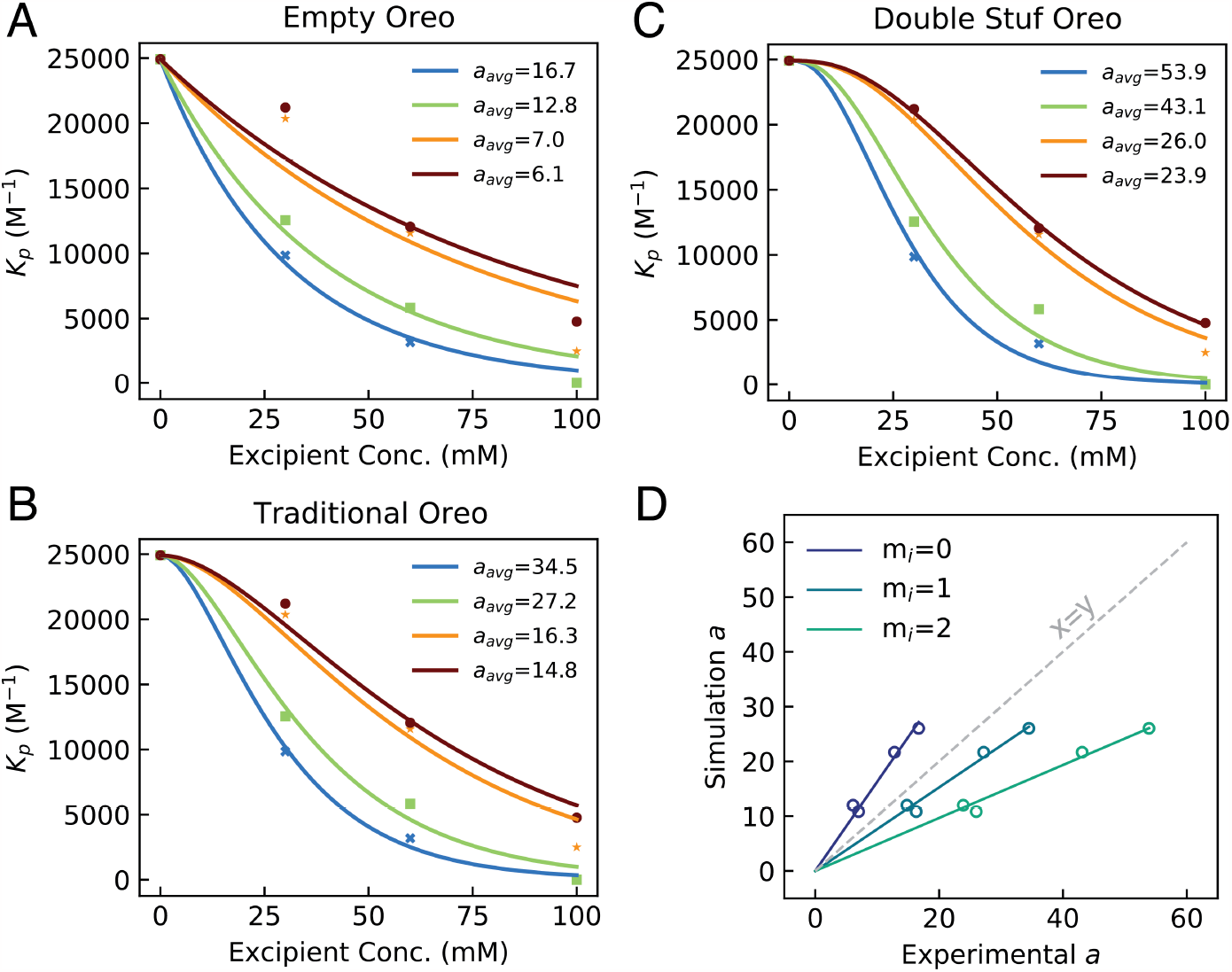
Fitting experimental data to the *Oreo* models, and comparing with predictions from simulations. A-C) The different *Oreo* models each fit experimental data reasonably well. Fitted parameters are included in the legend. D) The correlation between *a*_*avg*_ calculated from simulations, and the same calculated from interpreting experimental viscosity curves. We find a linear relationship that is dependent on the treatment of excipient molecules allowed inside the binding interface.

This approach is mostly successful in rank-ordering the effectiveness of each excipient at disrupting MEDI-4212 interactions, finding that Arg *>* Lys *>* Asp ≈ Glu. in agreement with experimental results. This indicates that arginine is the best at disrupting PPI, and will be most effective as a viscosity-lowering excipient, even at relatively low concentrations.

While lysine, aspartate and glutamate also work as viscosity reducers, higher concentrations are required to give the same effect as a lower concentration of arginine.

## Conclusions

We develop here a simple approach for computing the degree to which small-molecule excipients alter protein-protein binding affinities. We first use ClusPro and MD simulations to identify binding sites in the protein-protein interface, and estimate ligand affinities for those sites. Next, because excipients are relatively weak binders, we can use a binding polynomial to enumerate the binding modes and estimate the weakening of protein-protein affinity. We test this on a set of data for a monoclonal antibody, MedImmune mAb 4212 for the reduction of solution viscosities by four different excipient types. We find semi-quantitative agreement with experimental results. ^26^ While the set of data we test is rather small, the study is limited by the availability of studies in the literature that contain both viscosity data in the presence of multiple excipients, as well as sequence or structural data on the mAbs themselves. In future work, we hope to utilize larger data sets and refine the selection of criteria used to make predictions, and hope to demonstrate the transferability of this approach to many proteins and excipient pairings. Given this, we hope this approach will prove useful to predicting effects of excipients on mAb solutions, as well as other proteins with different solution behaviors.

## Code Available

Codes have been made available on Github which are useful for analyzing serial MD simulations of proteins and excipient molecules, and fitting to the binomial theory developed in this work.

## Acknowledgement

We thank Barbara Hribar-Lee for help with the Wertheim modeling and Pradipta Bandyopadhyay and Dima Kozakov for helpful discussions. This work was funded by the National Institutes of General Medical Sciences award 3RM1GM13513604S1.

## Supporting information for

### Supporting Text

#### Section S1: ClusPro docking identifies binding interfaces of Fab and Fc fragments of MEDI-4212

As a test case, we consider the MedImmune IgG1 mAb (MEDI-4212) that is studied in several previous papers [Arora, 2016; Hu, 2020; Cloutier,2019]. MEDI-4212 was shown to interact with both Fab and Fc fragments. The binding interfaces were identified using hydrogen-deuterium exchange mass spectrometry (HDX-MS) experiments [Arora, 2016], and show interfaces that are oppositely charged in the Fab and Fc regions, indicating that interactions are likely predominantly cross interactions between Fab and Fc fragments (Fig. S1A). We used the ClusPro rigid protein docking webserver [Kozakov, 2013; Kozakov, 2017; Vajda, 2017; Desta, 2020] to identify likely bound configurations of a single Fab fragment, and a single Fc fragment (Fig. S1B,C). Since HDX-MS yields information on the amino acid sequence, and which residues are buried, there is no one bound structure available, however, we can compare results from ClusPro to the sequences identified by these experiments by calculating the buried surface area of each amino acid in the sequence (Fig. S1B). Two of the identified conformations match all four of the identified regions of the sequence (Fig. S1C). We consider any residue that has a reduced solvent accessible surface area (SASA) by at least 4 Å^2^ to be a “buried’’ residue, and thus, part of the protein-protein interaction (PPI) interface in a given model.

We note that the two configurations that best match the experimental data are not ranked the highest (most likely) by ClusPro. Thus, in other cases where HDX-MS data is not available, further refinement or assessment must be done to determine which models are most reliable, such as refinement using molecular dynamics and enhanced sampling [Brini, 2019]. In this case, the first five predicted models capture at least two of the four identified sub-sequences, so an alternative approach may be to analyze excipient interactions with surfaces identified by multiple different models. We also note that even in the best models, the HDX-MS-identified regions are not the only parts of the protein that are buried. This is likely due to the limitation of the HDX-experiments, and flexibility of the proteins and the interface.

#### Section S2: Simulation details & analysis

Simulations were all conducted using the OpenMM software package [Eastman, 2017], using Langevin dynamics set to 300K, and a Monte Carlo barostat set to 1 bar. All simulations were carried out for 100 ns simulation time.

To analyze the simulations, we calculated the binding polynomial as a histogram of how many excipient molecules were within a certain cutoff distance (i.e. 4 Å) of any residue that was identified as part of the PPI interface in ClusPro model 3. The normalized histograms can then be fit to eq. 5. Since we conducted simulations a three different concentrations, we fit the model to all three histograms for a given excipient, defining the total error function as the root of the mean squared error:

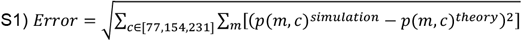

Note that in this optimization, we use all three concentrations together as the binding polynomials are concentration-dependent. For the second summation, we use all values of m where the probability from the simulation is non-zero.

A list of all simulations conducted for this work can be found in Table S1. If we ignore the protein molecule, each simulation is roughly 70, 140 or 210 mM of excipient. However, considering the large size of the protein in comparison with the box, we take into account the effect of volume excluded by the protein molecule. The volume fraction taken up by protein in the simulations is ∼0.1. Thus we divide each excipient concentration by 0.9 to account for this excluded volume.

#### Section S3: Wertheim’s thermodynamic perturbation theory for 7-bead mAb model

As in previous work [Kastelic, 2018], we use Wertheim’s thermodynamic perturbation theory to fit viscosity curves from the literature and extract interaction energies from the model. We model interactions between mAb fragments using a square well potential with a width of ω=0.18 nm, and a well depth of ε. Using integral equation theory, we solve for the fraction of bound Fab sites (*X*_*A*_ and *X*_*B*_) and the fraction of bound Fc sites (*X*_*C*_).

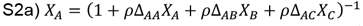

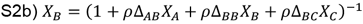

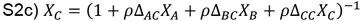

Where ρ is the protein concentration, and

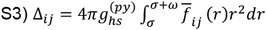

And 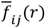 is the orientation average of the Mayer function for interactions between the interacting sites, and accounting for the flexibility in the 7-bead antibody model:

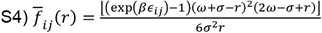

For the head-tail model of mAb interactions, all values of *ε*are set to zero except for interactions between the Fab and Fc fragments: *∈*_*AC*_ *= ∈*_*BC*_ ≠ 0. The values of ε reported the main text correspond to the values of *∈*_*AC*_ and *∈*_*BC*_ here.

We then calculate the cluster size distribution of the mAb solution based on the number of bound Fab and Fc sites, and polymer physics. For the head-tail model, we calculate the number distribution, H(n,c_P_) as:

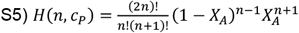

And the mass-weighted distribution is:

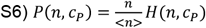

Note that the analytical solution for the average cluster size, <n> is simply equal to 1/X_C_.

Finally, the viscosity of the solution normalized to the low-concentration viscosity, η_0_,can be found using:

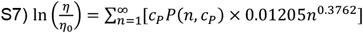

#### Section S4: Binding polynomial with non-equivalent excipient binding sites

Here justify the mathematical simplification that all binding sites are equivalent, having the same affinity for binding an excipient molecule (i.e. *K*_*e*,1_ *= K*_*e*,2_ *=*… *= K*_*e,N*_).

Perhaps a more realistic scenario would be that there is some distribution of strong and weak binding sites. Thus we would have ***K***_***e***_ *=* [***K***_***e***,**1**_, ***K***_***e***,**2**_, … ***K***_***e***,***N***_] where the values in *K*_*e*_ are drawn from a probability distribution. Given this construction, we can modify the binding polynomial to explicitly account for N unique binding sites each with their own unique affinity for an excipient molecule:

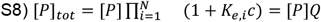

This defines a new excipient-binding partition function, *Q =* (1 + *K*_*e*,1_*c*)(1 + *K*_*e*,2_*c*)… (1 + *K*_*e,N*_), which can be easily applied to the empty Oreo model as well:

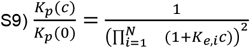

To demonstrate that the single binding site affinity assumption is reasonable, we compare two different cases with numbers close to what we are working with on MEDI-4212. The first case, α, has N binding sites of equivalent affinity 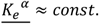, and the second case, β, also has N binding sites, but they are of different affinities, 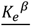and are pulled from an exponential distribution *f*(*x*, 1/*const*.). After generating the affinities from the exponential distribution, the single affinity for case α was set to the average affinity of sites in case β. We calculate the excipient concentration dependent protein binding affinity (*K*_*p*_(*c*)) using the empty Oreo model for both cases, and show they are nearly identical. We tested for two values of N at the high and low ends of what we might consider within the Oreo models to ensure that for low values of N, the statistical fluctuations do not result in significant deviations of the single-affinity model (Fig. S2).

#### Section S5: Definition of protein-protein binding interface for analysis

We defined the binding interface in the main text as any atom on any amino acid residue that is buried within the ClusPro-predicted bound conformation. To do this, we calculated the solvent accessible surface area (SASA) of each fragment in isolation, and then in the docked pose, and subtracted the difference:

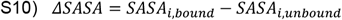

The per-residue ΔSASA is shown for all top 10 bound conformations from ClusPro in Fig. S1B. To translate this to a list of residues in the binding interface, we set a cutoff value of 2, 4 or 6 Å^2^. Thus any residue with Δ*SASA ≤ cutoff*is considered part of the protein-protein binding interface. The definition of a “buried” residue has an influence on the number of residues counted in the bound interface, as does the bound conformation that is used to make the ΔSASA calculation. Since both 3 and 4 match well with experimental HDX-MS data, we consider both cases, and three cutoffs to see how the extracted parameters differ.

In all cases, we optimized the model to fit the *a* and *N* parameters and calculate *K*_*e*_ *= a*/*N*. Since the *N* parameter has a minimal effect on the quality of the fit, we constrained it to be within reasonable boundaries (10 *≤ N ≤* 100). We show the calculated *a* values for both Fab and Fc from model 3 and model 4, and for all three definitions of the protein-protein interface in Fig. S4-6. We also tabulate the fitted parameters *K*_*e*_, *N*, and *a* in Tables S2-4. We note that regardless of the definition used for the interface, the qualitative comparison between excipients remains the same.

#### Section S6: Deriving a multi-excipient model and using it to ignore buffer

Since histidine buffer is present in all cases (at ∼10 mM) for the tested data, we need to consider that the histidine molecules in the solution may also have an effect on the protein-protein interactions. We thus derive a multi-excipient model that treats each excipient orthogonally, and has broader applications outside this work, perhaps to aid in development of multi-excipient formulations.

For each excipient, we can describe its binding polynomial using a few parameters, namely *K*_*e*_, *N*, and *a*. So if we have two excipients in the system, we need two values of each parameter, one for each excipient. As an example case, let’s use Arginine. For all cases of MEDI-4212 in solution with arginine, we have both arginine and histidine present in the solution. For ease, we will modify the empty Oreo model to account for a second excipient type.

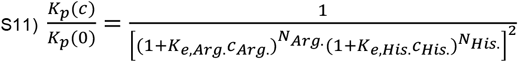

Within this model, the partition function of a single protein binding two types of excipient is generalized as *Q =* (1 + *K*_*e*,1_*c*_1_)^*N*1^ (1 + *K*_*e*,2_*c*_2_)^*N*2^. This can be further generalized for any number of excipient molecules as:

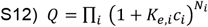

We now consider how the concentration of histidine is held constant for all experiments, thus the second term of the partition function, (1 + *K*_*e,His*._*c*_*His*._)^*N>His*.^ is constant. If we multiply both sides of eq. S11 by the square of this, we obtain:

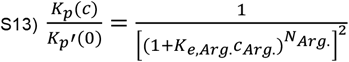

where *K*_*p*_′(0) *= K*_*p*_(0) × (1 + *K*_*e,His*._*c*_*His*._)^2*N>His*.^ is the protein-protein binding affinity at zero excipient with 10 mM histidine buffer accounted for.

### Supporting Figures

**Figure S1:**
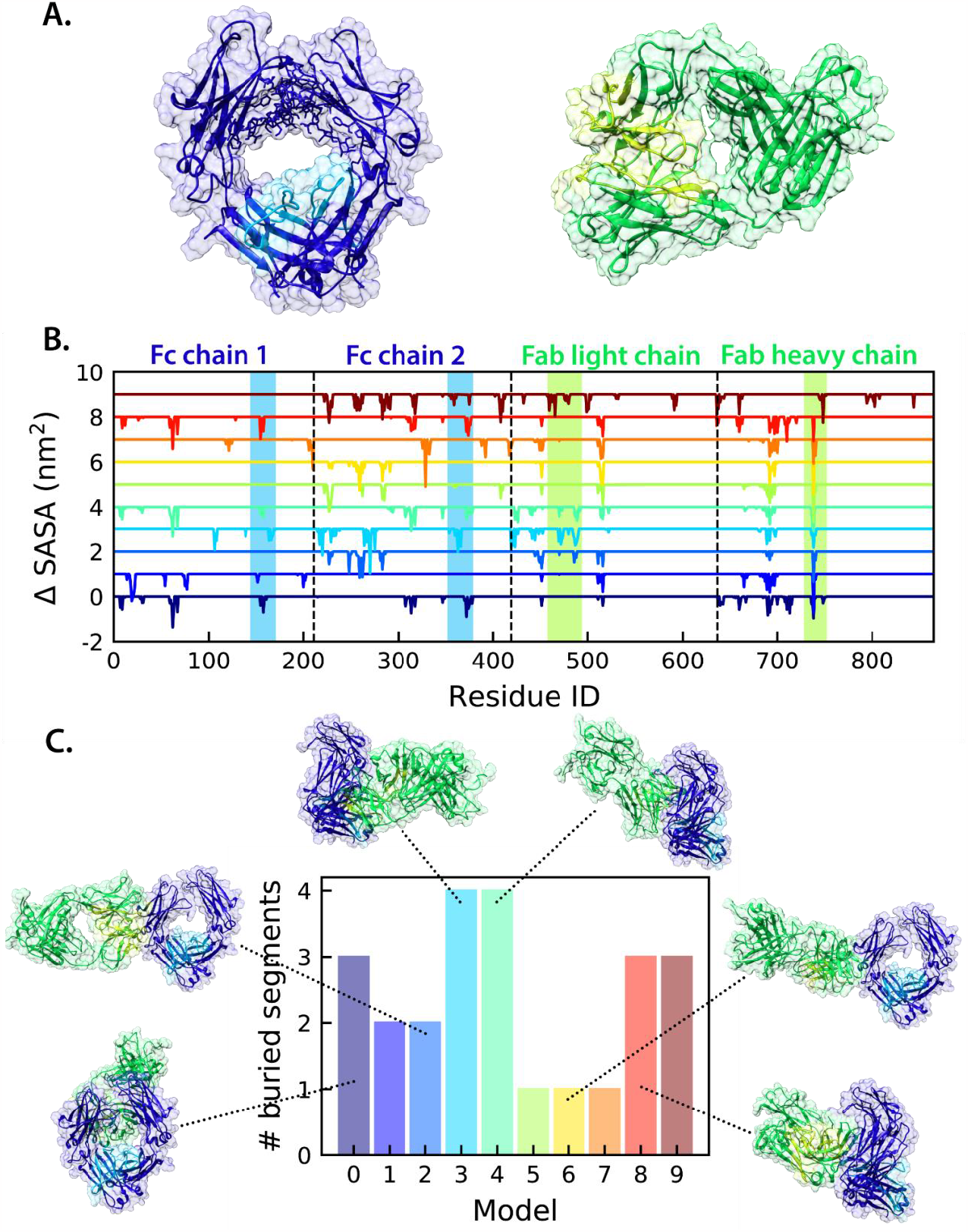
Docking identifies possible bound configurations in agreement with experimental data. **A)** Structures of Fc and Fab fragments of MEDI-4212 with interaction-prone regions from HDX-MS data shown in highlighted colors. **B)** Buried solvent accessible surface area (SASA) of each amino acid in the interacting mAb fragments for each of the top 10 identified bound conformations. Each ClusPro model is pushed up by +1 to aid in visualization. Segments that were identified from HDX-MS experiments are highlighted in blue (Fc) and lime green (Fab). **C)** The number of HDX-MS-identifed segments that have at least one amino acid buried in each of the top 10 ClusPro models. Selected models are visualized.

**Figure S2:**
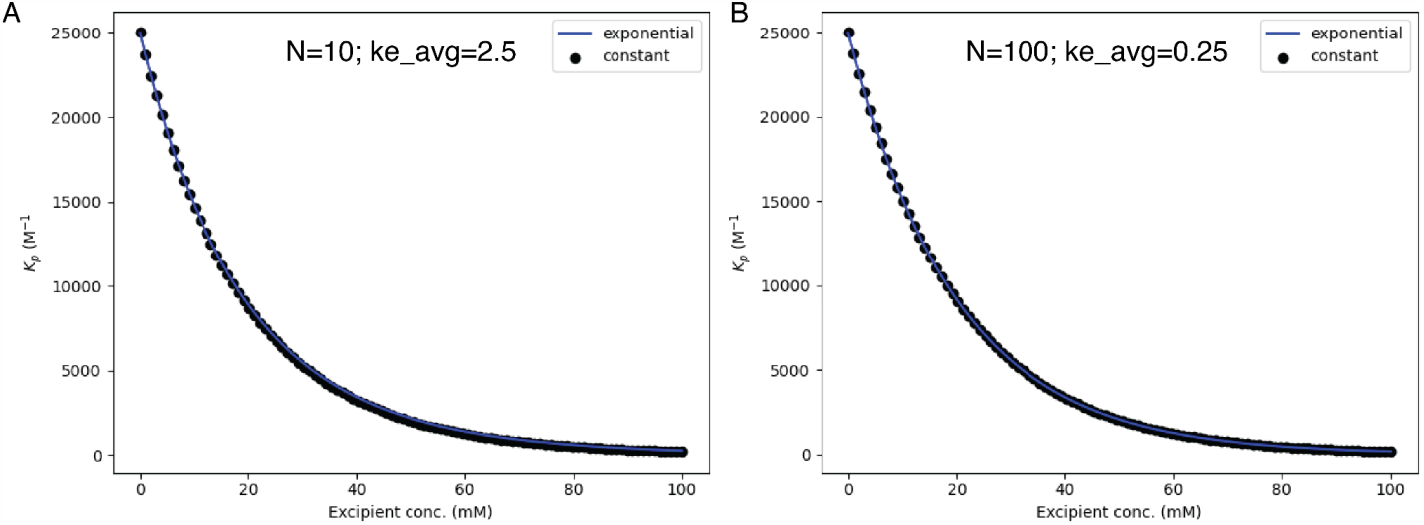
Using multi-affinity model and single-affinity model give nearly identical predictions of protein-protein binding affinity within the Empty Oreo model, even at small numbers of binding sites as low as 10.

**Figure S3:**
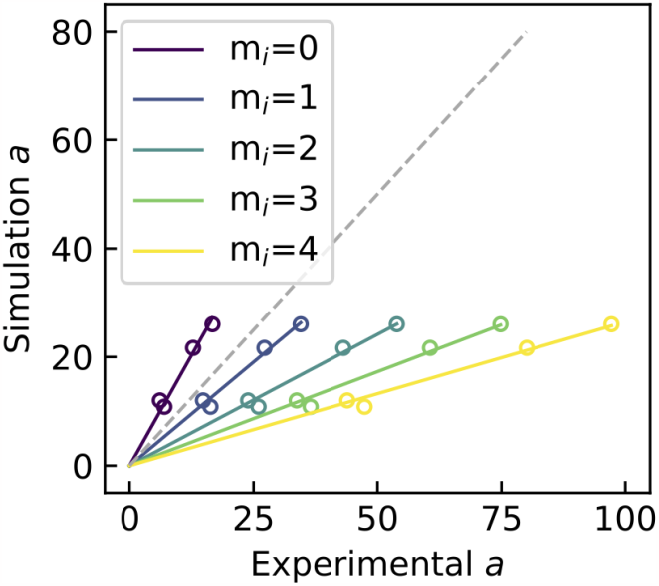
Higher order Oreo models do not improve prediction of excipient viscosity-reducing effects. The best predictor appears to be around 0-1 excipient molecules accommodated within the PPI interface.

**Figure S4:**
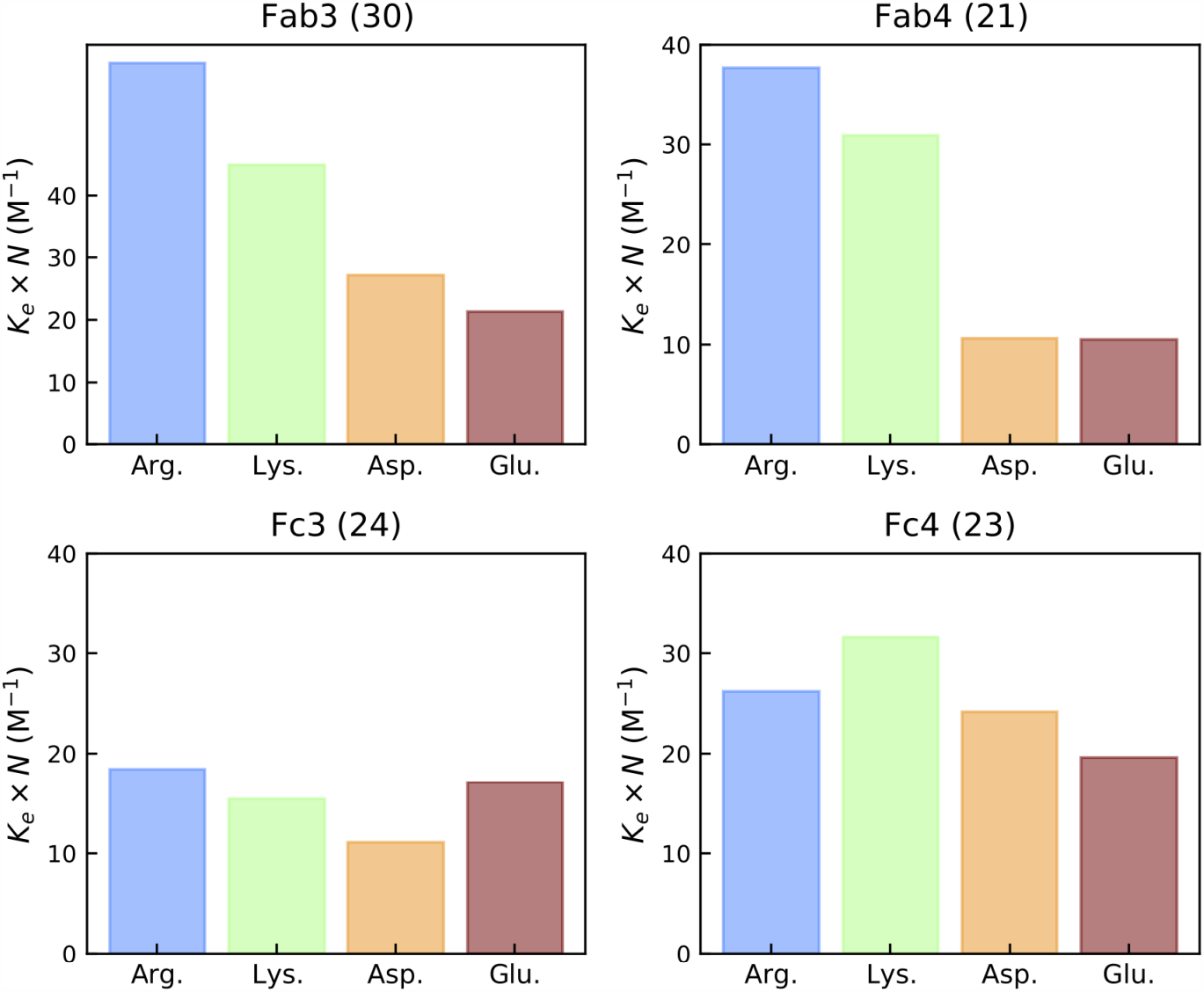
Combined *a* parameter for fitting the binomials to data from simulations gathered using the 2 Å^2^ cutoff to define the protein-protein interaction interface. The number of amino acid residues considered “buried” are included in the subplot titles in parentheses.

**Figure S5:**
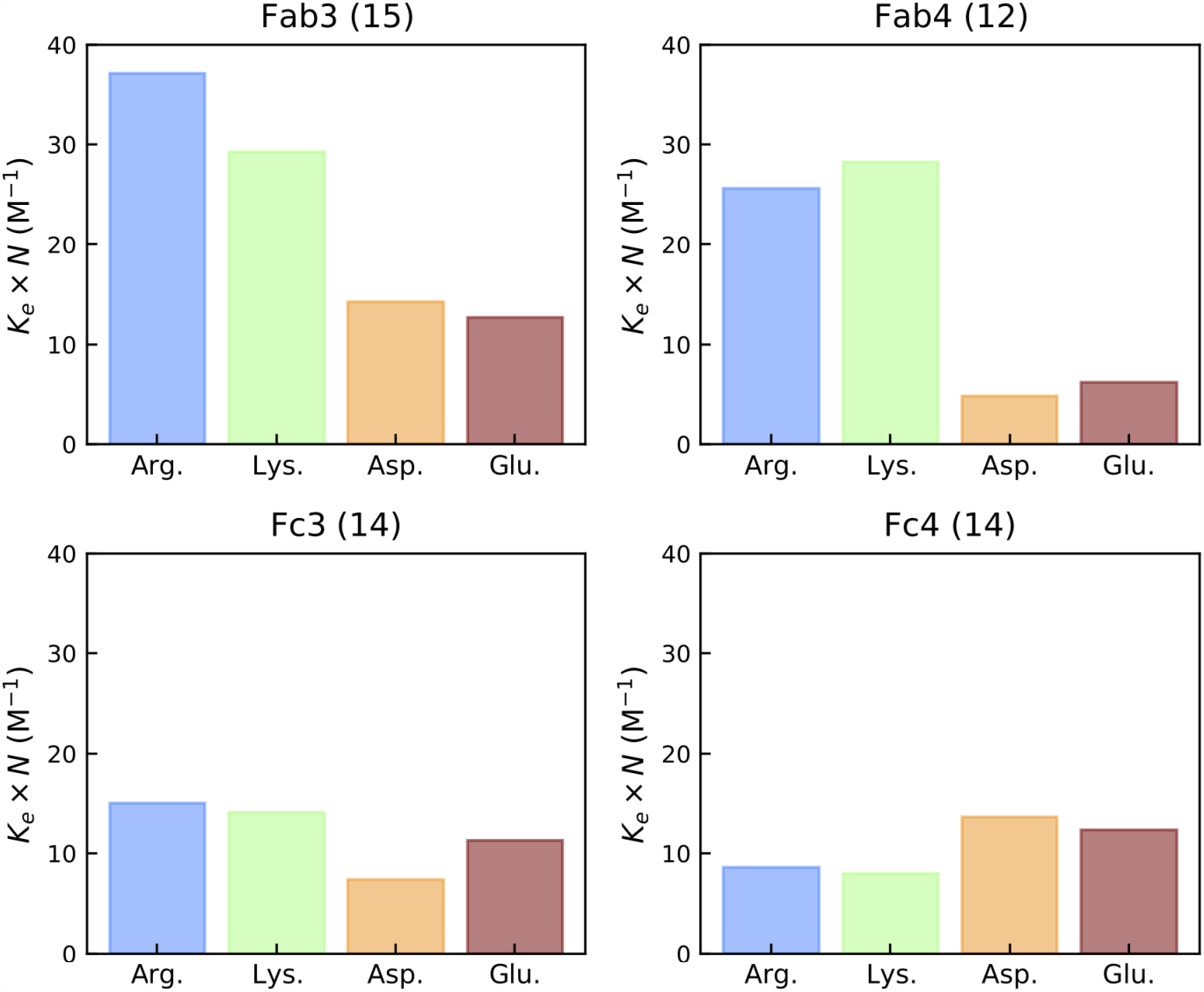
Combined *a* parameter for fitting the binomials to data from simulations gathered using the 4 Å^2^ cutoff to define the protein-protein interaction interface. The number of amino acid residues considered “buried” are included in the subplot titles in parentheses.

**Figure S6:**
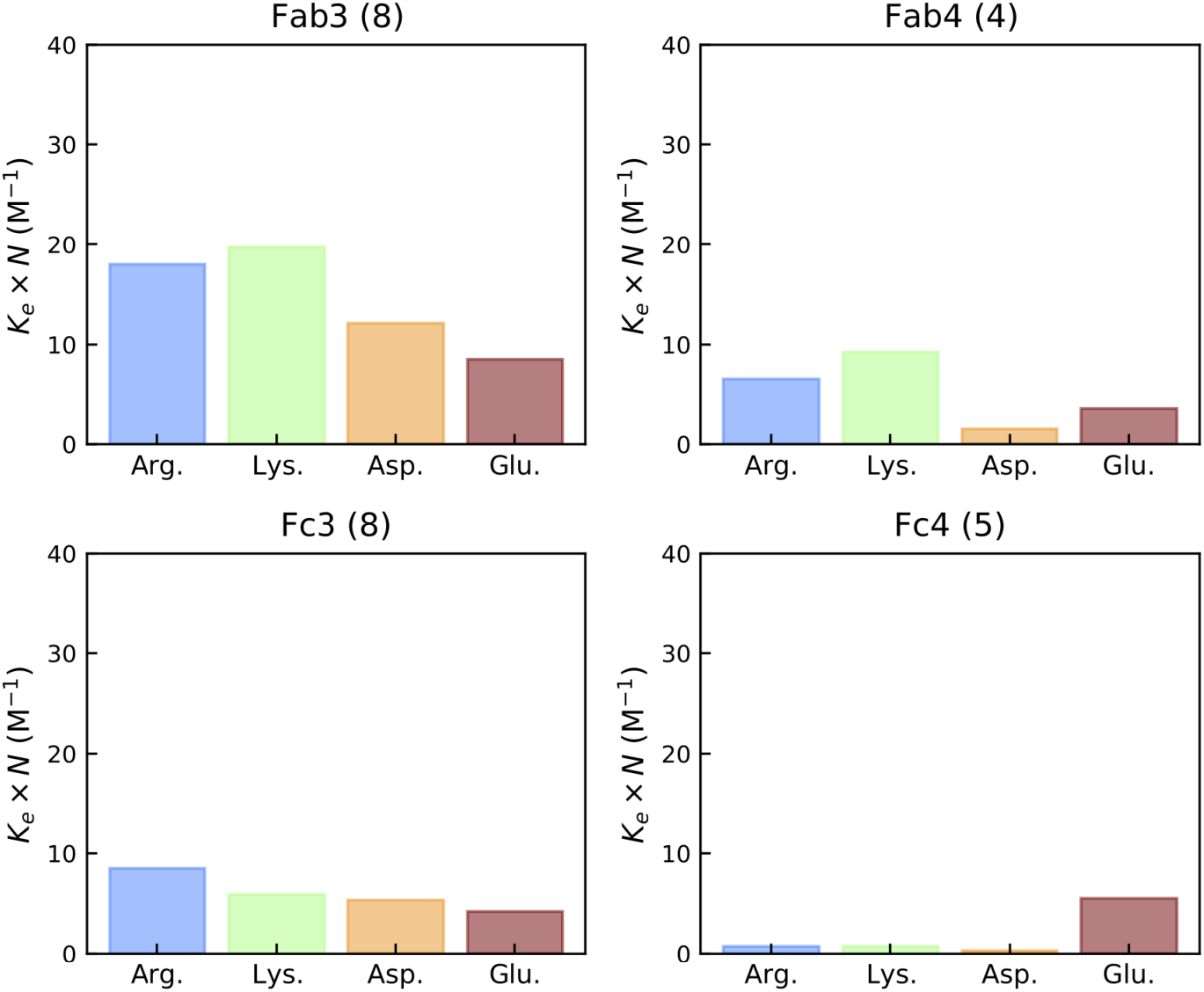
Combined *a* parameter for fitting the binomials to data from simulations gathered using the 6 Å^2^ cutoff to define the protein-protein interaction interface. The number of amino acid residues considered “buried” are included in the subplot titles in parentheses.

**Figure S7:**
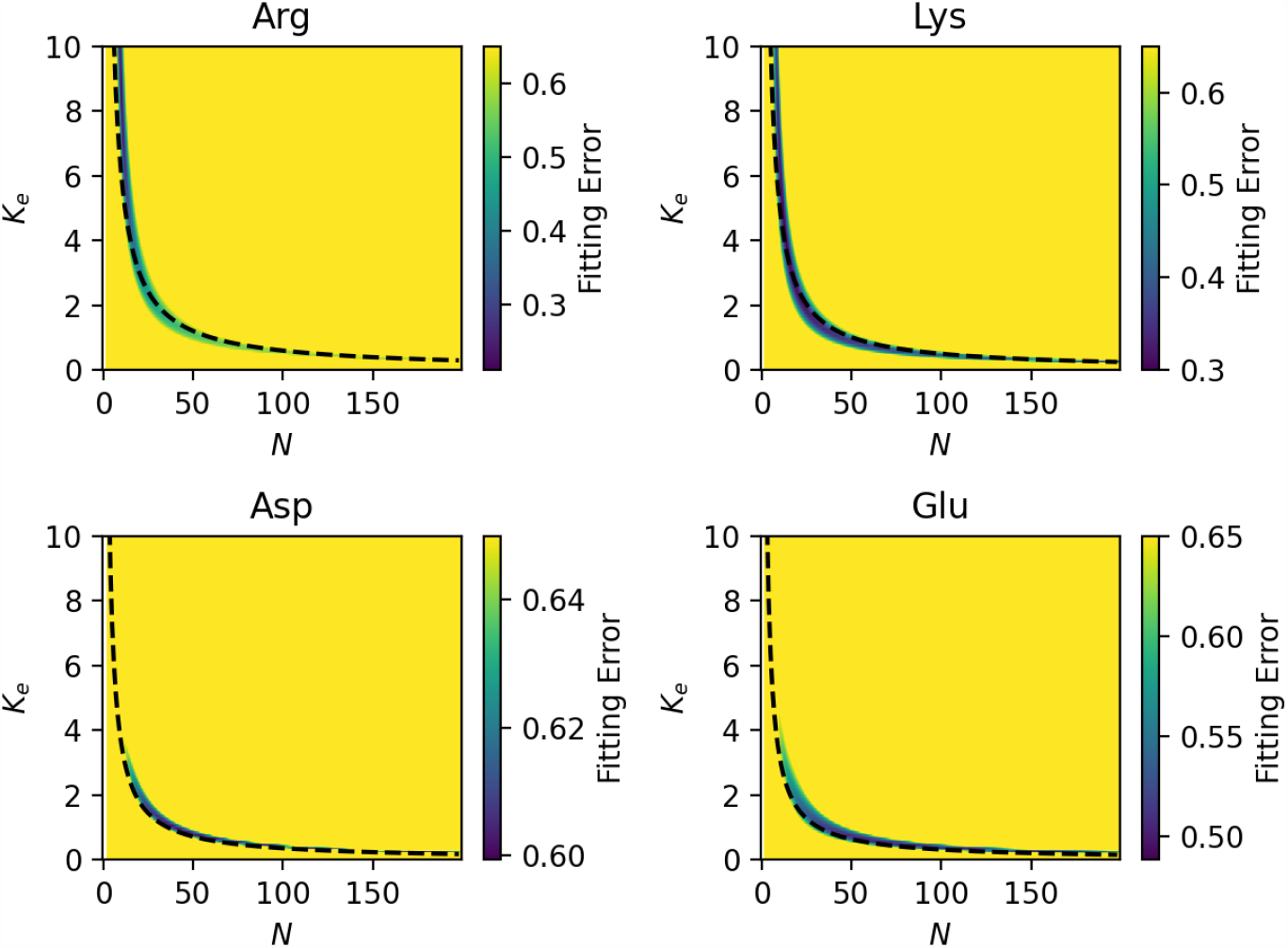
Fitting error is high for all combinations of *K*_*e*_ and *N* except for one region that follows roughly an inverse curve. This region of low error indicates an optimal fit of the binomial to the data from simulations. Dashed lines represent the relationship *K*_*e*_ *= a*/*N*.

**Figure S8:**
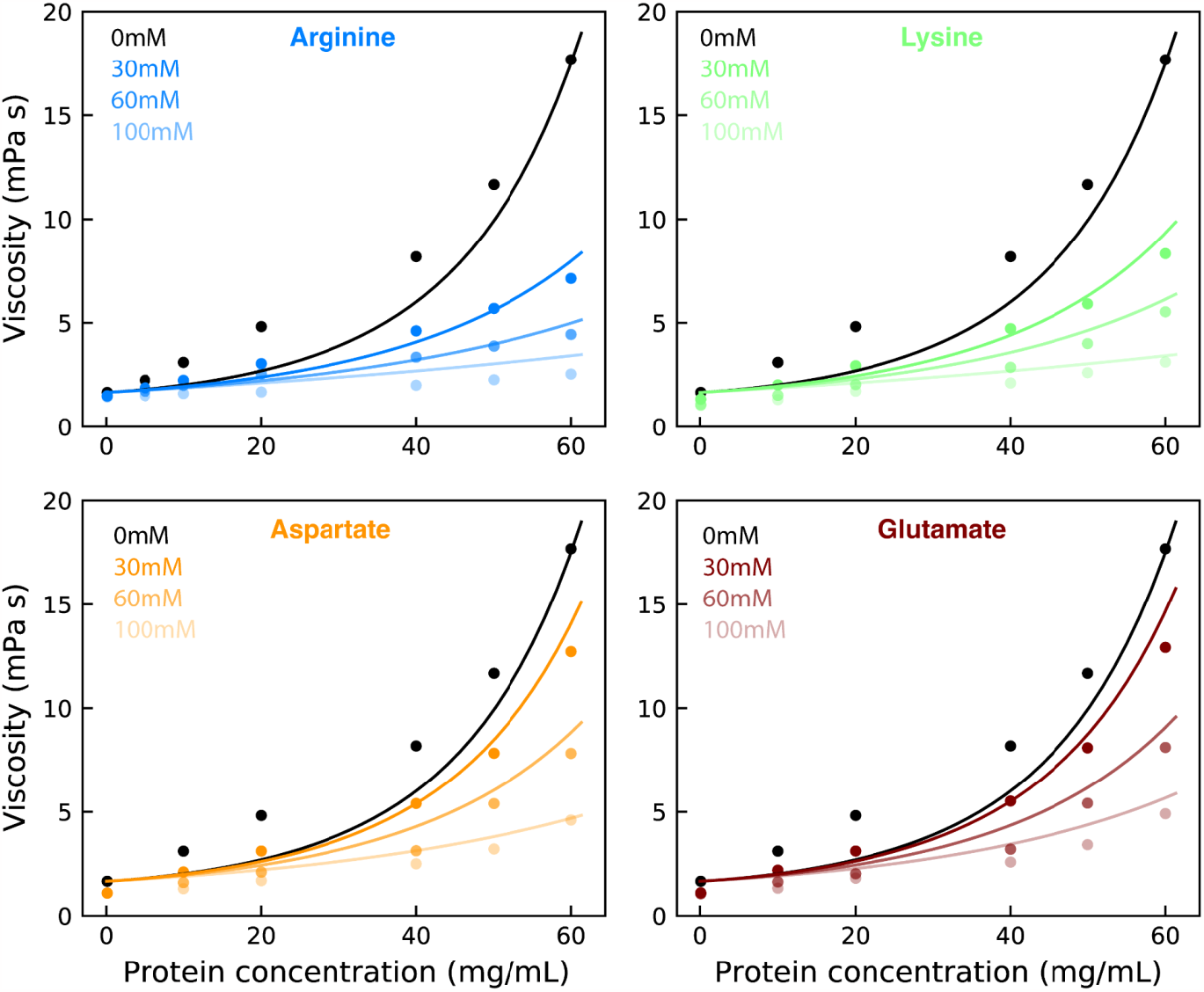
Data points from experimental work are shown as points, and fits using Wertheim’s thermodynamic perturbation theory are solid lines. We find that the theory fits all data sets reasonably well using the head-tail interaction model.

**Figure S9:**
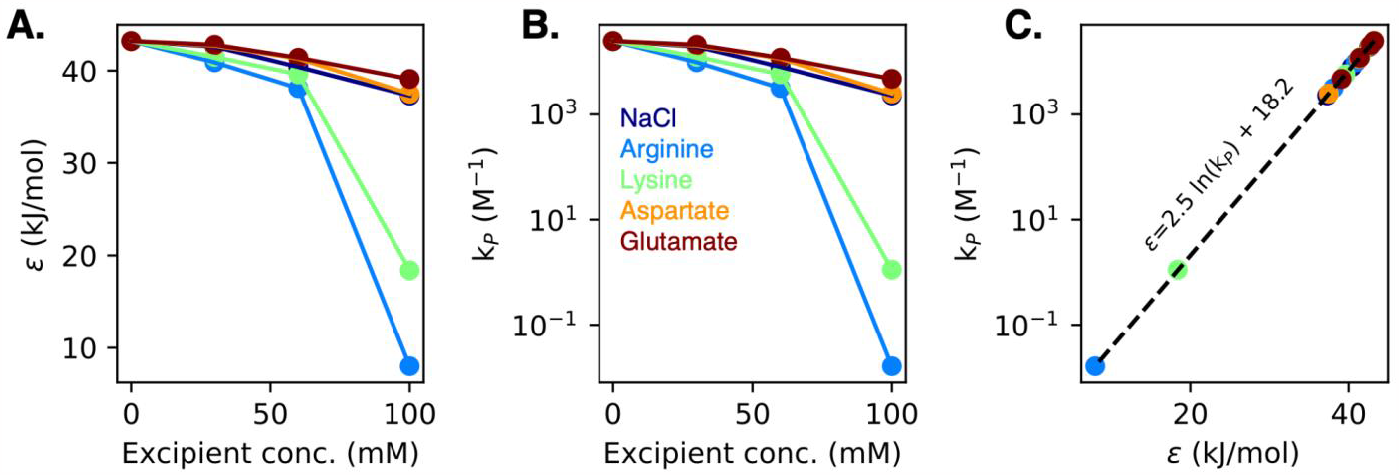
Comparison of calculated values from Wertheim’s thermodynamic perturbation theory. **A)** Well depth, ε, for square well potential as a function of excipient concentration. **B)** Association constant, K_p_ calculated from population of monomers and dimers at very low concentrations. Note that this is plotted on a log scale here for comparison with ε, and is plotted on a linear scale in the main text. **C)** The two values are related using the thermodynamic relationship with an intercept value of 18.2 which likely responds to an entropic cost of binding and a slope of 2.5 which is equal to RT.

**Figure S10:**
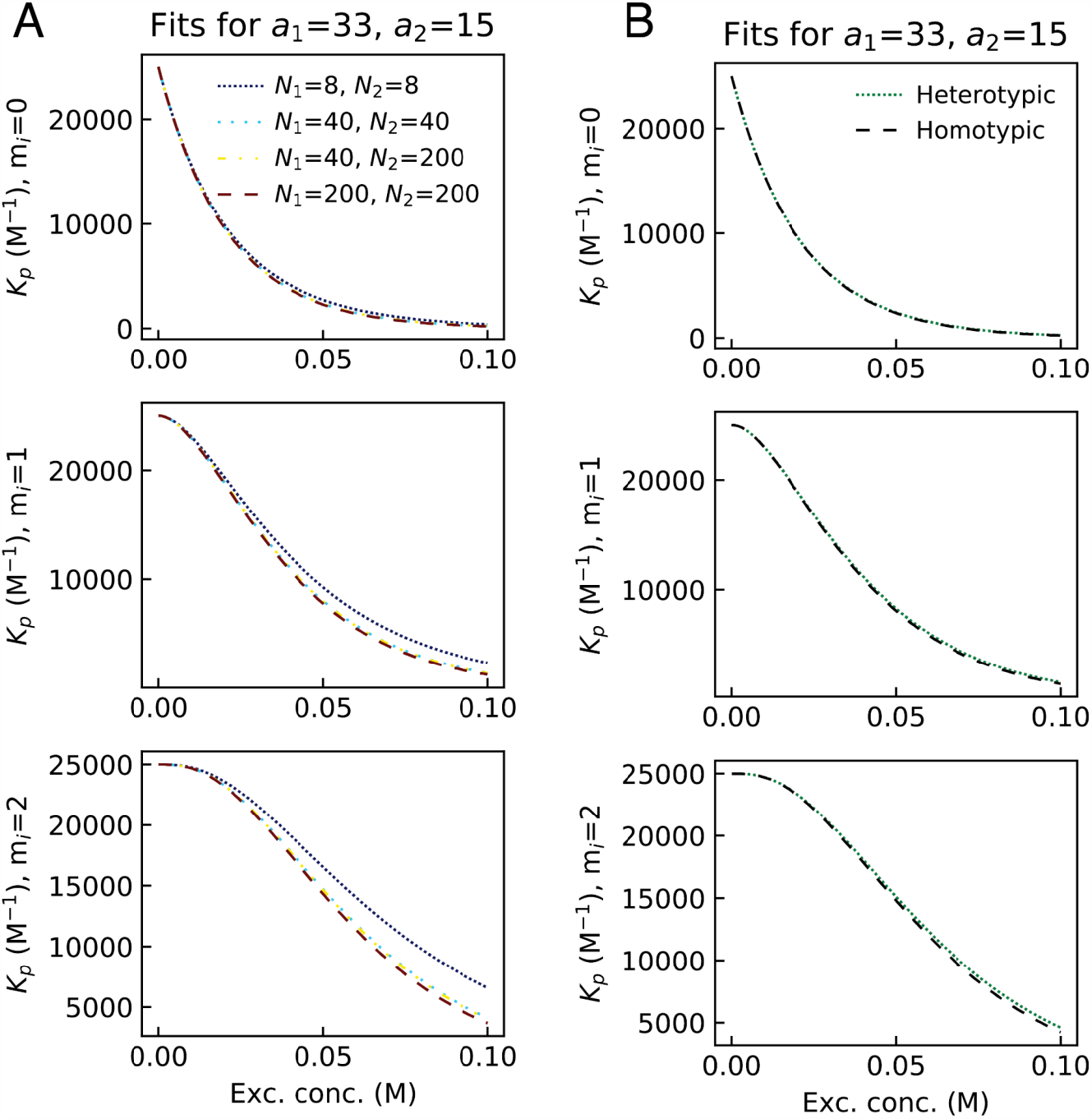
Sensitivity of Empty, traditional, and double stuffed Oreo models to different simplifications. **A)** Different values of N have a very small effect on the predictions of K_p_ from all of the Oreo models. Having very low values, however, is most likely to result in some deviation from the other values. This is most pronounced in the double-stuffed model. **B)** All three models are very well-captured by a homotypic model that assumes an average of the a parameters as well as an average of the N parameters. For the heterotypic models, we use N_1_=20 and N_2_=40, and for homotypic, N_avg_=30.

### Supporting Tables

**Table S1:** List of simulations conducted for this work.

| Fragment | Excipient | #<sub>exc</sub> (conc. mM) | Fragment | Excipient | #<sub>exc</sub> (conc. mM) |
| --- | --- | --- | --- | --- | --- |
| Fab | Arginine | 25 (77) | Fc | Arginine | 25 (77) |
| Fab | Arginine | 50 (154) | Fc | Arginine | 50 (154) |
| Fab | Arginine | 75 (231) | Fc | Arginine | 75 (231) |
| Fab | Lysine | 25 (77) | Fc | Lysine | 25 (77) |
| Fab | Lysine | 50 (154) | Fc | Lysine | 50 (154) |
| Fab | Lysine | 75 (231) | Fc | Lysine | 75 (231) |
| Fab | Aspartate | 25 (77) | Fc | Aspartate | 25 (77) |
| Fab | Aspartate | 50 (154) | Fc | Aspartate | 50 (154) |
| Fab | Aspartate | 75 (231) | Fc | Aspartate | 75 (231) |
| Fab | Glutamate | 25 (77) | Fc | Glutamate | 25 (77) |
| Fab | Glutamate | 50 (154) | Fc | Glutamate | 50 (154) |
| Fab | Glutamate | 75 (231) | Fc | Glutamate | 75 (231) |

**Table S2:** Tabulated values from fits to binding binomial from simulations using the 2 Å^2^ cutoff to define the protein-protein interaction interface.

|  | Fab model 3 (30 res) |  |  | Fab model 4 (21 res) |  |  | Fc model 3 (24 res) |  |  | Fc model 4 (23 res) |  |  |
| --- | --- | --- | --- | --- | --- | --- | --- | --- | --- | --- | --- | --- |
| Excipient | N | Ke | a | N | Ke | a | N | Ke | a | N | Ke | a |
| Arg. | 13.2 | 4.634 | 61.171 | 9.1 | 4.135 | 37.627 | 10 | 1.844 | 18.44 | 10 | 2.62 | 26.202 |
| Lys. | 14.6 | 3.076 | 44.905 | 14.9 | 2.074 | 30.903 | 10 | 1.543 | 15.434 | 36.2 | 0.873 | 31.597 |
| Asp. | 100 | 0.272 | 27.154 | 100 | 0.106 | 10.625 | 10 | 1.11 | 11.103 | 100 | 0.242 | 24.17 |
| Glu. | 100 | 0.213 | 21.319 | 100 | 0.105 | 10.507 | 10 | 1.71 | 17.104 | 10 | 1.962 | 19.613 |

**Table S3:** Tabulated values from fits to binding binomial from simulations using the 4 Å^2^ cutoff to define the protein-protein interaction interface. The values highlighted in green are those used in the main text.

|  | Fab model 3 (15 res) |  |  | Fab model 4 (12 res) |  |  | Fc model 3 (14 res) |  |  | Fc model 4 (14 res) |  |  |
| --- | --- | --- | --- | --- | --- | --- | --- | --- | --- | --- | --- | --- |
| Excipient | N | Ke | a | N | Ke | a | N | Ke | a | N | Ke | a |
| Arg. | 10 | 3.708 | 37.075 | 11.6 | 2.209 | 25.627 | 10 | 1.5 | 15.002 | 10 | 0.861 | 8.614 |
| Lys. | 13.9 | 2.105 | 29.266 | 17.9 | 1.579 | 28.256 | 10 | 1.408 | 14.083 | 10 | 0.8 | 7.999 |
| Asp. | 24.6 | 0.58 | 14.264 | 100 | 0.049 | 4.879 | 10 | 0.74 | 7.396 | 10 | 1.362 | 13.622 |
| Glu. | 100 | 0.127 | 12.703 | 100 | 0.063 | 6.254 | 10 | 1.129 | 11.291 | 10 | 1.234 | 12.335 |

**Table S4:** Tabulated values from fits to binding binomial from simulations using the 6 Å^2^ cutoff to define the protein-protein interaction interface.

|  | Fab model 3 (8 res) |  |  | Fab model 4 (5 res) |  |  | Fc model 3 (8 res) |  |  | Fc model 4 (4 res) |  |  |
| --- | --- | --- | --- | --- | --- | --- | --- | --- | --- | --- | --- | --- |
| Excipient | N | Ke | a | N | Ke | a | N | Ke | a | N | Ke | a |
| Arg. | 10.4 | 1.73 | 17.996 | 10.0 | 0.658 | 6.578 | 100.0 | 0.085 | 8.507 | 10.0 | 0.074 | 0.741 |
| Lys. | 10.0 | 1.971 | 19.714 | 10.0 | 0.926 | 9.256 | 10.0 | 0.591 | 5.91 | 24.4 | 0.03 | 0.728 |
| Asp. | 21.0 | 0.578 | 12.134 | 10.0 | 0.153 | 1.527 | 10.0 | 0.536 | 5.358 | 24.8 | 0.013 | 0.32 |
| Glu. | 100 | 0.085 | 8.537 | 10.0 | 0.364 | 3.639 | 100.0 | 0.042 | 4.215 | 10.0 | 0.552 | 5.523 |

**Table S5:** Buried residues included in each of the Fab and Fc models with different SASA cutoffs. Note that numbering of residues corresponds with sequences listed in Table S6. Model 3+4 indicates residues that are present in either model 3 or 4 at the given cutoff definition.

| Model | Cutoff def | Fab Residues | Fc Residues |
| --- | --- | --- | --- |
| 3 | 2Å<sup>2</sup> | **Light:** Q1, V3, T5, P8, T23, S25, S26, G30, D34, Y51, D52, F54, N55, R56, D62, S67, K68, S69, G70, T71, S72, **Heavy:** D54, N55, E57, E62, K63, G101, W103, I104, K105 | **Fc1:** R107, A108, P109, I140, G165, Y167, F168, M169, **Fc2:** D12, M15, C24, V27, V29, Y59, N60, W62, R64, V65, V66, P70, K155, E158, V160, L161 |
| 3 | 4Å<sup>2</sup> | **Light:** Q1, V3, T5, T23, S26, Y51, F54, N55, K68, S69, G70, T71, **Heavy:** D54, E57, W103 | **Fc1:** R107, P109, G165, Y167, **Fc2:** D12, M15, C24, Y59, W62, R64, V65, P70, E158, L161 |
| 3 | 6Å<sup>2</sup> | **Light:** Q1, V3, T5, F54, S69, G70, **Heavy:** D54, W103 | **Fc1:** R107, **Fc2:** D12, M15, Y59, V65, P70, E158, L161 |
| 4 | 2Å<sup>2</sup> | **Light:** T5, Q6, P7, P8, S21, T23, D52, S69, G70, T71, S72, P96, L98, T104, **Heavy:** D54, N55, E57, F59, E62, K102, W103 | **Fc1:** P8, I10, E31, N60, S61, W62, L63, R64, A68, N156, P159, **Fc2:** K89, K103, G104, P109, Q110, V111, V113, V142, Y167, Y170, S171, L173 |
| 4 | 4Å<sup>2</sup> | **Light:** T5, P8, D52, S69, T71, P96, L98, **Heavy:** D54, N55, E57, K102, W103 | **Fc1:** P8, I10, S61, W62, L63, A68, P159, **Fc2:** K103, P109, Q110, V111, V113, V142, Y167 |
| 4 | 6Å<sup>2</sup> | **Light:** P8, T71, **Heavy:** E57, K102, W103 | **Fc1:** L63, A68, **Fc2:** P109, V113 |
| 3+4 | 2Å<sup>2</sup> | **Light:** Q1, V3, T5, Q6, P7, P8, S21, T23, S25, S26, G30, D34, Y51, D52, F54, N55, R56, D62, S67, K68, S69, G70, T71, S72, P96, L98, T104, **Heavy:** D54, N55, E57, F59, E62, K63, G101, K102, W103, I104, K105 | **Fc1:** P8, I10, E31, N60, S61, W62, L63, R64, A68, R107, A108, P109, I140, N156, P159, G165, Y167, F168, M169, **Fc2:** D12, M15, C24, V27, V29, Y59, N60, W62, R64, V65, V66, P70, K89, K103, G104, P109, Q110, V111, V113, V142, K155, E158, V160, L161, Y167, Y170, S171, L173 |
| 3+4 | 4Å<sup>2</sup> | **Light:** Q1, V3, T5, P8, T23, S26, Y51, D52, F54, N55, K68, S69, G70, T71, P96, L98, **Heavy:** D54, N55, E57, K102, W103 | **Fc1:** P8, I10, S61, W62, L63, A68, R107, P109, P159, G165, Y167, **Fc2:** D12, M15, C24, Y59, W62, R64, V65, P70, K103, P109, Q110, V111, V113, V142, E158, L161, Y167 |
| 3+4 | 6Å<sup>2</sup> | **Light:** Q1, V3, T5, P8, F54, S69, G70, T71, **Heavy:** D54, E57, K102, W103 | **Fc1:** L63, A68, R107, **Fc2:** D12, M15, Y59, V65, P70, P109, V113, E158, L161 |

**Table S6:**
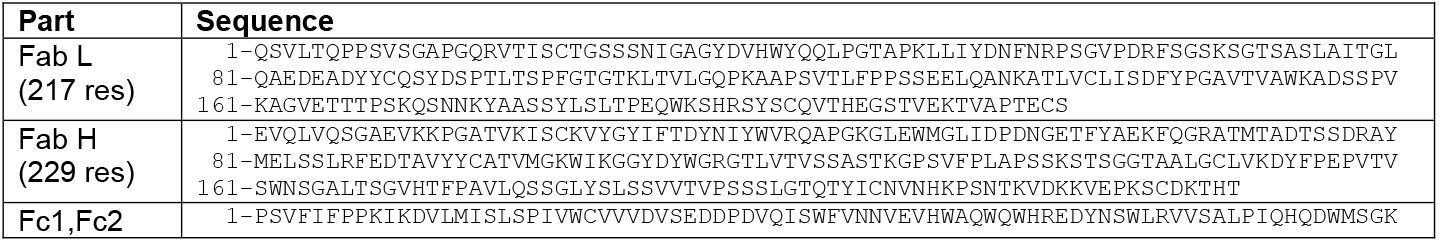
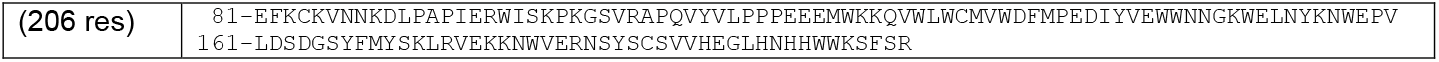
Sequences of Fab and Fc fragments used in simulations. Both chains in Fc fragment are identical

## References

(1) Klinge, S.; Woolford Jr, J. L. Ribosome assembly coming into focus. Nature reviews Molecular cell biology 2019, 20, 116–131.

(2) Prasad, B. V.; Hardy, M. E.; Dokland, T.; Bella, J.; Rossmann, M. G.; Estes, M. K. X-ray crystallographic structure of the Norwalk virus capsid. Science 1999, 286, 287–290.

(3) Brangwynne, C. P.; Tompa, P.; Pappu, R. V. Polymer physics of intracellular phase transitions. Nature Physics 2015, 11, 899–904.

(4) Dignon, G. L.; Best, R. B.; Mittal, J. Biomolecular phase separation: from molecular driving forces to macroscopic properties. Annual review of physical chemistry 2020, 71, 53–75.

(5) Banani, S. F.; Lee, H. O.; Hyman, A. A.; Rosen, M. K. Biomolecular condensates: organizers of cellular biochemistry. Nature reviews Molecular cell biology 2017, 18, 285–298.

(6) Boeynaems, S.; Alberti, S.; Fawzi, N. L.; Mittag, T.; Polymenidou, M.; Rousseau, F.; Schymkowitz, J.; Shorter, J.; Wolozin, B.; Van Den Bosch, L.; others Protein phase separation: a new phase in cell biology. Trends in cell biology 2018, 28, 420–435.

(7) Schmit, J. D.; Ghosh, K.; Dill, K. What drives amyloid molecules to assemble into oligomers and fibrils? Biophysical journal 2011, 100, 450–458.

(8) Chiti, F.; Dobson, C. M. Protein misfolding, amyloid formation, and human disease: a summary of progress over the last decade. Annual review of biochemistry 2017, 86, 27–68.

(9) Shire, S. J.; Shahrokh, Z.; Liu, J. Challenges in the Development of High Protein Concentration Formulations. Journal of Pharmaceutical Science 2004, 1390–1402.

(10) Shire, S. J.; Shahrokh, Z.; Liu, J. Challenges in the development of high protein con-centration formulations. Current trends in monoclonal antibody development and manufacturing 2010, 131–147.

(11) Barnett, G. V.; Qi, W.; Amin, S.; Lewis, E. N.; Roberts, C. J. Aggregate structure, morphology and the effect of aggregation mechanisms on viscosity at elevated protein concentrations. Biophysical Chemistry 2015, 207, 21–29.

(12) Tomar, D. S.; Kumar, S.; Singh, S. K.; Goswami, S.; Li, L. Molecular basis of high viscosity in concentrated antibody solutions: strategies for high concentration drug product development. MAbs 2016, 8, 216–228.

(13) Barnett, G. V.; Razinkov, V. I.; Kerwin, B. A.; Blake, S.; Qi, W.; Curtis, R. A.; Roberts, C. J. Osmolyte effects on monoclonal antibody stability and concentrationdependent protein interactions with water and common osmolytes. The Journal of Physical Chemistry B 2016, 120, 3318–3330.

(14) Braun, A.; Kwee, L.; Labow, M. A.; Alsenz, J. Protein aggregates seem to play a key role among the parameters influencing the antigenicity of interferon alpha (IFN-α) in normal and transgenic mice. Pharmaceutical research 1997, 14, 1472–1478.

(15) Wang, W.; Ohtake, S. Science and art of protein formulation development. International journal of pharmaceutics 2019, 568, 118505.

(16) Du, Q.; Damschroder, M.; Pabst, T. M.; Hunter, A. K.; Wang, W. K.; Luo, H. Process optimization and protein engineering mitigated manufacturing challenges of a monoclonal antibody with liquid-liquid phase separation issue by disrupting inter-molecule electrostatic interactions. MAbs 2019, 11, 789–802.

(17) Kastelic, M.; Dill, K. A.; Kalyuzhnyi, Y. V.; Vlachy, V. Controlling the viscosities of antibody solutions through control of their binding sites. Journal of molecular liquids 2018, 270, 234–242.

(18) Kastelic, M.; Vlachy, V. Theory for the liquid–liquid phase separation in aqueous antibody solutions. The Journal of Physical Chemistry B 2018, 122, 5400–5408.

(19) Ramallo, N.; Paudel, S.; Schmit, J. Cluster formation and entanglement in the rheology of antibody solutions. The Journal of Physical Chemistry B 2019, 123, 3916–3923.

(20) von Bülow, S.; Siggel, M.; Linke, M.; Hummer, G. Dynamic cluster formation determines viscosity and diffusion in dense protein solutions. Proceedings of the National Academy of Sciences 2019, 116, 9843–9852.

(21) Zheng, W.; Dignon, G. L.; Jovic, N.; Xu, X.; Regy, R. M.; Fawzi, N. L.; Kim, Y. C.; Best, R. B.; Mittal, J. Molecular details of protein condensates probed by microsecond long atomistic simulations. The Journal of Physical Chemistry B 2020, 124, 11671–11679.

(22) Cloutier, T.; Sudrik, C.; Mody, N.; Sathish, H. A.; Trout, B. L. Molecular computations of preferential interaction coefficients of IgG1 monoclonal antibodies with sorbitol, sucrose, and trehalose and the impact of these excipients on aggregation and viscosity. Molecular pharmaceutics 2019, 16, 3657–3664.

(23) Jo, S.; Xu, A.; Curtis, J. E.; Somani, S.; MacKerell Jr, A. D. Computational Characterization of Antibody–Excipient Interactions for Rational Excipient Selection Using the Site Identification by Ligand Competitive Saturation-Biologics Approach. Molecular pharmaceutics 2020, 17, 4323–4333.

(24) Fry, D. C. Protein–protein interactions as targets for small molecule drug discovery. Peptide Science: Original Research on Biomolecules 2006, 84, 535–552.

(25) Arora, J.; Hu, Y.; Esfandiary, R.; Sathish, H. A.; Bishop, S. M.; Joshi, S. B.; Middaugh, C. R.; Volkin, D. B.; Weis, D. D. Charge-mediated Fab-Fc interactions in an IgG1 antibody induce reversible self-association, cluster formation, and elevated viscosity. MAbs 2016, 8, 1561–1574.

(26) Hu, Y.; Arora, J.; Joshi, S. B.; Esfandiary, R.; Middaugh, C. R.; Weis, D. D.; Volkin, D. B. Characterization of excipient effects on reversible self-association, backbone flexibility, and solution properties of an IgG1 monoclonal antibody at high con-centrations: part 1. Journal of Pharmaceutical Sciences 2020, 109, 340–352.

(27) Cohen, E. S.; Dobson, C. L.; Käck, H.; Wang, B.; Sims, D. A.; Lloyd, C. O.; England, E.; Rees, D. G.; Guo, H.; Karagiannis, S. N.; others A novel IgE-neutralizing antibody for the treatment of severe uncontrolled asthma. MAbs 2014, 6, 755–763.

(28) Kirschner, K. N.; Yongye, A. B.; Tschampel, S. M.; González-Outeiriño, J.; Daniels, C. R.; Foley, B. L.; Woods, R. J. GLYCAM06: a generalizable biomolecular force field. Carbohydrates. Journal of computational chemistry 2008, 29, 622–655.

(29) Kozakov, D.; Hall, D. R.; Xia, B.; Porter, K. A.; Padhorny, D.; Yueh, C.; Beglov, D.; Vajda, S. The ClusPro web server for protein–protein docking. Nature protocols 2017, 12, 255–278.

(30) Kozakov, D.; Beglov, D.; Bohnuud, T.; Mottarella, S. E.; Xia, B.; Hall, D. R.; Vajda, S. How good is automated protein docking? Proteins: Structure, Function, and Bioinformatics 2013, 81, 2159–2166.

(31) Vajda, S.; Yueh, C.; Beglov, D.; Bohnuud, T.; Mottarella, S. E.; Xia, B.; Hall, D. R.; Kozakov, D. New additions to the C lus P ro server motivated by CAPRI. Proteins: Structure, Function, and Bioinformatics 2017, 85, 435–444.

(32) Desta, I. T.; Porter, K. A.; Xia, B.; Kozakov, D.; Vajda, S. Performance and its limits in rigid body protein-protein docking. Structure 2020, 28, 1071–1081.

(33) Eastman, P.; Swails, J.; Chodera, J. D.; McGibbon, R. T.; Zhao, Y.; Beauchamp, K. A.; Wang, L.-P.; Simmonett, A. C.; Harrigan, M. P.; Stern, C. D.; others OpenMM 7: Rapid development of high performance algorithms for molecular dynamics. PLoS computational biology 2017, 13, e1005659.

(34) Maier, J. A.; Martinez, C.; Kasavajhala, K.; Wickstrom, L.; Hauser, K. E.; Simmerling, C. ff14SB: improving the accuracy of protein side chain and backbone parameters from ff99SB. Journal of chemical theory and computation 2015, 11, 3696–3713.

(35) Jorgensen, W. L.; Chandrasekhar, J.; Madura, J. D.; Impey, R. W.; Klein, M. L. Comparison of simple potential functions for simulating liquid water. The Journal of chemical physics 1983, 79, 926–935.

(36) Wang, J.; Wolf, R. M.; Caldwell, J. W.; Kollman, P. A.; Case, D. A. Development and testing of a general amber force field. Journal of computational chemistry 2004, 25, 1157–1174.

(37) Hopkins, C. W.; Le Grand, S.; Walker, R. C.; Roitberg, A. E. Long-time-step molecular dynamics through hydrogen mass repartitioning. Journal of chemical theory and computation 2015, 11, 1864–1874.

(38) Flory, P. J. Principles of polymer chemistry ; Cornell university press, 1953.

(39) Aune, K. C.; Tanford, C. Thermodynamics of the denaturation of lysozyme by guanidine hydrochloride. II. Dependence on denaturant concentration at 25. Biochemistry 1969, 8, 4586–4590.

(40) Wyman, J.; Gill, S. J. Binding and linkage: functional chemistry of biological macromolecules; University Science Books, 1990.

(41) Dill, K.; Bromberg, S. Molecular driving forces: statistical thermodynamics in biology, chemistry, physics, and nanoscience; Garland Science, 2010.

(42) Dill, K.; Jernigan, R. L.; Bahar, I. Protein actions: Principles and modeling; Garland Science, 2017.

(43) Jumper, J.; Evans, R.; Pritzel, A.; Green, T.; Figurnov, M.; Ronneberger, O.; Tunyasuvunakool, K.; Bates, R.; Žídek, A.; Potapenko, A.; others Highly accurate protein structure prediction with AlphaFold. Nature 2021, 596, 583–589.

(44) Evans, R.; O’Neill, M.; Pritzel, A.; Antropova, N.; Senior, A.; Green, T.; Žídek, A.; Bates, R.; Blackwell, S.; Yim, J.; others Protein complex prediction with AlphaFold-Multimer. biorxiv 2021, 2021–10.

